# The Boltzmann distributions of folded molecular structures predict likely changes through random mutations

**DOI:** 10.1101/2023.02.22.529545

**Authors:** Nora S. Martin, Sebastian E. Ahnert

**Affiliations:** Theory of Condensed Matter Group, Cavendish Laboratory, University of Cambridge, JJ Thomson Avenue, Cambridge CB3 0HE, UK; Sainsbury Laboratory, University of Cambridge, Bateman Street, Cambridge CB2 1LR, UK; Rudolf Peierls Centre for Theoretical Physics, Beecroft Building, Parks Road, Oxford OX1 3PU, UK; Department of Chemical Engineering and Biotechnology, University of Cambridge, Philippa Fawcett Drive, Cambridge CB3 0AS, UK; The Alan Turing Institute, 96 Euston Road, London NW1 2DB, UK

## Abstract

New folded molecular structures can only evolve after arising through mutations. This aspect is modelled using genotype-phenotype (GP) maps, which connect sequence changes through mutations to changes in molecular structures. Previous work has shown that the likelihood of appearing through mutations can differ by orders of magnitude from structure to structure and that this can affect the outcomes of evolutionary processes. Thus, we focus on the phenotypic mutation probabilities *ϕ* _*qp*_, i.e. the likelihood that a random mutation changes structure *p* into structure *q*. For both RNA secondary structures and the HP protein model, we show that a simple biophysical principle can explain and predict how this likelihood depends on the new structure *q*: *ϕ* _*qp*_ is high if sequences that fold into *p* as the minimum-free-energy structure are likely to have *q* as an alternative structure with high Boltzmann frequency. This generalises the existing concept of plastogenetic congruence from individual sequences to the entire neutral spaces of structures. Our result helps us understand why some structural changes are more likely than others, can be used as a basis for estimating these likelihoods via sampling and makes a connection to alternative structures with high Boltzmann frequency, which could be relevant in evolutionary processes.

## I. INTRODUCTION

For a new molecular structure to evolve, it first has to appear through random mutations. This is not just a qualitative statement, but a quantitative one: if a specific structure appears sooner and more frequently in an evolutionary process, it has a higher chance of going into fixation within a given time frame [3–5]. This theoretical argument is supported by evolved RNA structures in databases, which tend to be structures that are likely to appear through random mutations [4, 6, 7]. Thus, the likelihood that a given structure will arise through random mutations is crucial for evolutionary processes.

Variation through random mutations can be modelled using a sequence-structure, or genotype-phenotype (GP) map, where random sequence mutations can be mapped to structural changes [8]. Due to the huge number of possible sequences (i.e. genotypes) and structures (i.e. phenotypes), this map is best studied computationally, for example using the ViennaRNA package for RNA folding [9] and the HP lattice model for protein folding [10]. Here, we focus on one key quantity: the phenotypic mutation probability *ϕ* _*qp*_, which describes how likely a random mutation on an initial structure *p* is to generate a specific structure *q* (as illustrated schematically in Fig. 1A). We follow the notation in [1, 3], but this quantity has been defined repeatedly [4, 11] due to its importance in evolutionary processes: in the biologically relevant case, where most structural changes are deleterious, a population will maintain its initial structure and only accumulate sequence changes that do not affect the folded structure. Thus the population is confined to the *neutral set* of the initial structure *p*, i.e. the set of sequences that fold into *p*. While most structural changes are strongly deleterious in this scenario, a small set of structural changes may be adaptive and so it is important to know how likely these structures will arise through random mutations. This is described by the *ϕ* _*qp*_ values [3] and since *ϕ* _*qp*_ values can vary over several orders of magnitude, the likelihood that a structure arises can differ from structure to structure by several orders of magnitude [3, 4].

**Figure 1.**
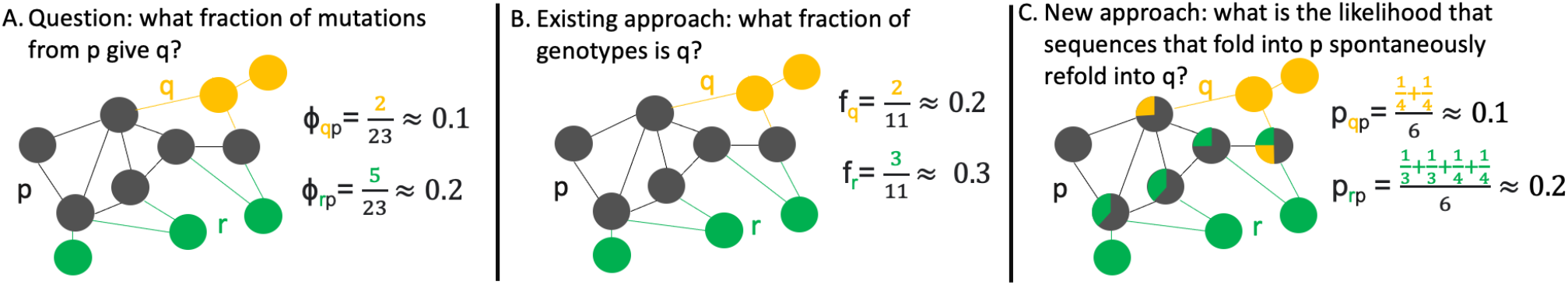
Schematic of *ϕ* _*qp*_ and the two approaches for approximating *ϕ* _*qp*_ differences: A) *ϕ* _*qp*_ is the fraction of all mutations from an initial phenotype (p, grey) to a new phenotype (q or r). Here, each node represents a genotype, its colour the corresponding phenotype and an edge represents a possible point mutation. We see that there are two different point mutations from *p* (grey) to *q* (yellow), out of a total of 23 mutations that start on the grey neutral set (mutations between two grey genotypes are counted twice since they could start on either genotype). This gives *ϕ* _*qp*_ = 2*/*23. The same procedure can be applied to compute *ϕ* _*rp*_. B) Existing approach [1]: the assumption is that the phenotypic frequency of *q* is a good indicator of *ϕ* _*qp*_. Since two out of eleven genotypes in the entire map corresponds to *q*, we have *f*_*q*_ = 3*/*11. C) New approach: each genotype that primarily folds into the initial phenotype *p* also folds into one or more suboptimal phenotypes with a Boltzmann frequency that is indicated by the pie charts (visualised as in [2]). Thus, we can compute the average Boltzmann frequency of *q* in the neutral set of *p* as *p*_*qp*_ = 1*/*12.

Because previous work has shown that *ϕ* _*qp*_ values are crucial for evolutionary processes, it is important to have a good understanding of which structures have high *ϕ* _*qp*_ values and why. As a first approximation it has been shown that the mutation probability *ϕ* _*qp*_ from *p* to *q* is proportional to the phenotypic frequency *f*_*q*_, which is the probability that an arbitrary sequence folds into *q* [1, 3](as illustrated schematically in Fig. 1B). However, this does not work well if the initial phenotype *p* has a low phenotypic frequency [1], and even for the highest-frequency phenotype the correlation was only found to be moderate in the HP protein model [1] (Spearman coeff. *ρ ≈* 0.5). Other work has focused exclusively on RNA structures and characterised what type of structural changes are likely to occur through mutations for RNA structures, finding that a new structure *q* is most likely to occur if it is similar to *p* and if the differences between *p* and *q* fall into specific classes (for example losses of stacks) [12]. Similarly, there have been several independent attempts to estimate the probability of protein structural changes that depend both on the initial and the new structure [13, 14]. However, all these methods only apply to a particular type of molecule - RNA or proteins at a specific level of coarse-graining - and cannot be generalised. In very recent work, an information-theoretic viewpoint has been used to predict *ϕ* _*qp*_ differences [15].

Here we use a biophysical perspective to gain a deeper understanding of *ϕ* _*qp*_ differences: in the GP map, each sequence is assumed to fold into a single structure, its minimum-free-energy (mfe) structure. However, other suboptimal structures can exist in addition to that structure [16], and these form the Boltzmann ensemble of that sequence. A principle termed *plastogenetic congruence* postulates that the suboptimal structures in this Boltzmann ensemble can indicate for a specific *sequence*, which structural changes are likely after mutations [17]. This has been shown not only for RNA [17], but also for proteins [18, 19]. This principle can be generalised from sequences to structures as follows: the mean probability that a sequence that primarily fold into *p* occupies structure *q* as a suboptimal state could be an indicator of the mutation probability *ϕ* _*qp*_ (as illustrated schematically in Fig. 1C). While the mutation probabilities for a specific sequence are known to differ markedly from the average mutation probabilities for the corresponding structure [1] and a similar finding exist for the set of energetically low-lying suboptimal structures [20], our previous work on insertion/deletion mutations in RNA [21] nevertheless shows that the concept of plastogenetic congruence can be generalised from sequences to structures, at least for a coarse-grained method of RNA structure prediction. This generalisation from sequences to structures is also conjectured in Ancel and Fontana’s original paper on plastogenetic congruence [17], but not tested quantitatively. Here, we test this approach more systematically for two classic molecular GP map models, RNA secondary structures and the HP protein model. We find that the new approach outperforms the existing frequency-based approach in predicting the most likely structural changes, i.e. the ones with the highest *ϕ* _*qp*_ values, and that is more accurate than approaches based on information theory or simple structural similarity measures. Our approach thus provides a biophysical way of understanding why some structural changes are likely to occur through mutations.

## II. RESULTS

### A. *ϕ*_*qp*_ and Boltzmann frequencies

We use two models in this analysis: the ViennaRNA model of RNA secondary structures and the HP lattice model of protein folding. First, we compare our new approach of approximating mutation probabilities *ϕ* _*qp*_ to the existing frequency-based approach for one specific choice of *p* as an example: the existing approach [1, 3] simply assumes that the probability *ϕ* _*qp*_ of obtaining *q* through mutations from *p* is described by the phenotypic frequency *f*_*q*_ of *q*, which is the probability that an arbitrary sequence folds into *q* (Fig. 2A&D for RNA and proteins respectively). The second approach is our new hypothesis: we plot *ϕ* _*qp*_ against a biophysical quantity, which we denote by *p*_*qp*_ (Fig. 2B&E for RNA and proteins respectively). This *p*_*qp*_ is defined as the mean Boltzmann frequency of *q* for a sequence that has *p* as its minimum-free-energy structure. Thus it is a measure of how likely temporary switches to *q* are to occur without the presence of mutations. We find that both approaches capture some of the trends in the *ϕ* _*qp*_ data, but the correlation is much clearer in the second case.

**Figure 2.**
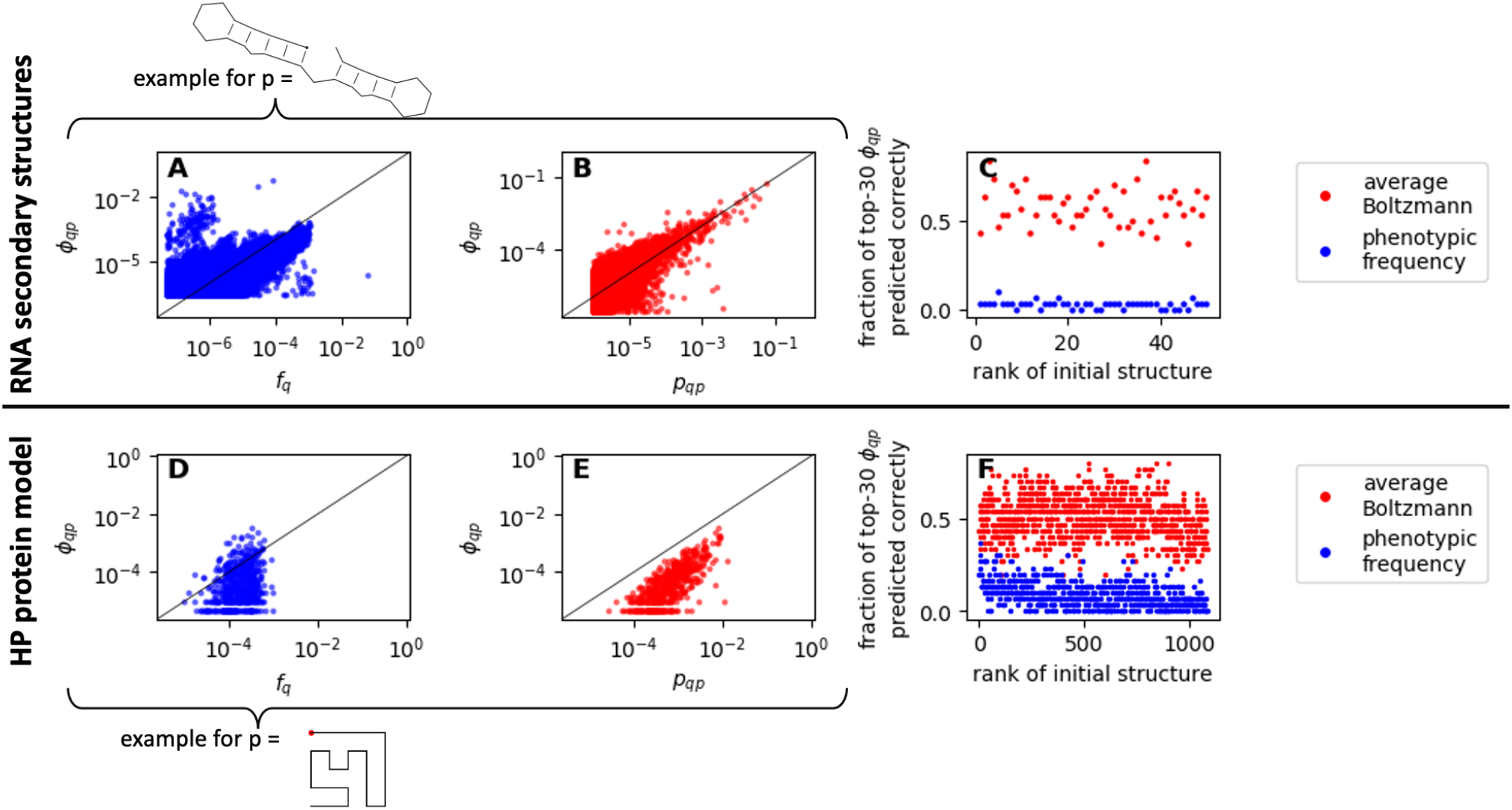
Comparison of the old and new approach for RNA secondary structures (A-C) and HP protein structures (D-F): (A) Old approach for one initial RNA structure *p* as an example: the *ϕ* _*qp*_ values for mutating from this initial structure *p* to a new structure *q* are plotted against the phenotypic frequency *f*_*q*_ of the new structure *q*. (B) New approach for the same initial RNA structure *p*: the *ϕ* _*qp*_ values for this initial structure are plotted against the Boltzmann frequency *p*_*qp*_ of the new structure *q* averaged over sequences in the neutral set of the initial structure *p*. (C) The analysis is repeated for 50 different initial structures *p*, each denoted by a unique rank (x-axis), and the approaches are compared by how many of the structures with the thirty highest *ϕ* _*qp*_ values they predict correctly (y-axis). This metric is 0% in the example in (A) and 67% in the example in (B). The rank denoting each structure *p* (x-axis) was defined, such that out of the 50 structures in the sample, the structure with the lowest phenotypic frequency has the highest rank. (D-F) Same for the HP protein model - here all 1081 structures that fold as mfe structures are included in F and the initial structure used in D & E is sketched below (the red dot indicates the start position of the sequence on the lattice). The fraction of structures with the thirty highest *ϕ* _*qp*_ values that are predicted correctly is 17% for (D) and 50% for E. Due to the log-scale, zero values are not shown in plots A, B, D & E and the black lines indicate *x* = *y*.

In order to quantify the performance of both approaches, we evaluate, how many of the structures with the thirty highest *ϕ* _*qp*_ values are predicted correctly by each of these methods. This method is relatively insensitive to sampling errors in our calculations, which are likely to predominantly affect low values of *f*_*q*_, *ϕ* _*qp*_ and *p*_*qp*_. We find that, in our RNA example, the new approach correctly predicts 20 out of the top-30 *ϕ* _*qp*_-structures (i.e. ≈ 67%), compared to the existing approach, which does not make any correct predictions. Thus, the accuracy is high even though we use two separate sequence samples to estimate *ϕ* _*qp*_ and *p*_*qp*_ (overlap *<* 0.01% on average for all initial structures *p* in this analysis) to ensure that we do not simply capture the known [17] correspondence between Boltzmann ensembles and mutational effects for specific *sequences*, but instead capturing the average properties of specific *structures*. In the protein example, the new approach also outperforms the existing approach, with 15 structures (i.e. 50%) predicted correctly, compared to five structures (i.e. ≈ 17%) in the existing approach. Unlike for the RNA example, sampling is only used for the *p*_*qp*_ values in the HP model. The advantage of this is that the data for the existing approach is exact in the HP model, but this also means that there is no clear separation between the sequences used to estimate *p*_*qp*_ and those used in the *ϕ* _*qp*_ calculations.

To conclude, neutral-set-averages of Boltzmann frequencies, *p*_*qp*_, are a good indicator of which structural transitions are likely. This conclusion generalises when we repeat the analysis for further initial structures *p*, including both structures with high and low phenotypic frequency (Fig. 2C&F for RNA and proteins respectively): our new approach is better at capturing the structural changes that are most likely to happen through random mutations. The full data for a range of structures is shown in section S3 in the SI. In addition, section S4 in the SI compares our results to approaches based on simple measures of structural similarity and to the recent algorithmic-information-theory-based approach from ref [15]. We find that our new approach outperforms all alternative approaches.

### B. Boltzmann frequencies and phenotypic frequencies

To better understand the neutral-set-averages of Boltzmann frequencies *p*_*qp*_, which are at the centre of our new approach, we also evaluated a closely related quantity, namely the averages of Boltzmann frequencies over random sequences. The data in Fig. 3 indicates that this quantity *p*_*q*_, the mean Boltzmann frequency of structure *q* over a random sequence sample, is correlated with the phenotypic frequency of that structure. This holds regardless of whether mfe structures are included in the averages, as in Fig. 3, or not, as shown in section S1 of the SI. This result agrees with existing computational data for the RNAshapes model (SI of ref [6]) and is consistent with our previous work showing correlations between various phenotypic frequency definitions in RNA [22]. It is also consistent with theoretical arguments [23] for a Boltzmann-like trend in phenotypic frequencies, where energetically unfavourable features are associated with an exponential decrease in phenotypic frequencies. However, not all assumptions behind this theoretical claim are met in our case (for example that the mean free energy of folding for a random sequence differs from structure to structure, which is not true in many compact HP models [24], and the assumption of constant sequence composition in the random energy model [25]). The fact that even the key assumption on free energy distributions is not met for the compact HP model may partially explain, why the correlation is less clear for the HP model.

**Figure 3.**
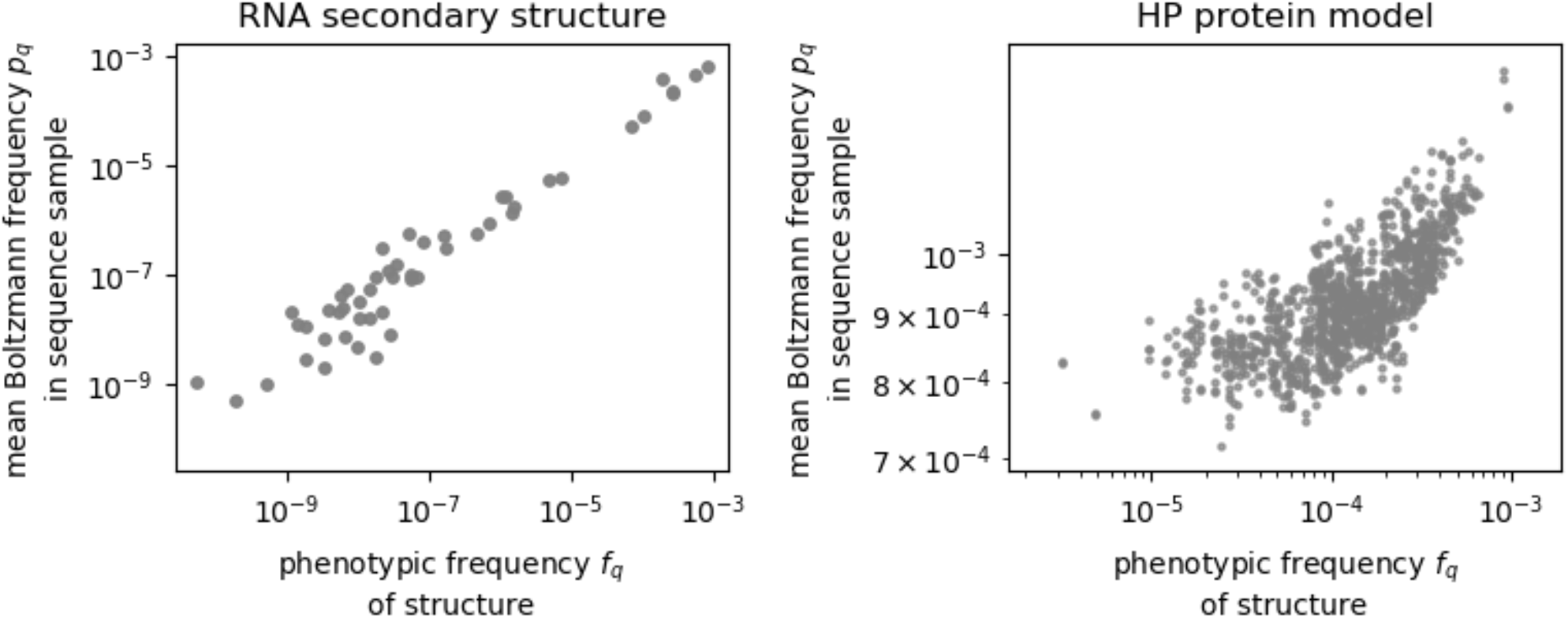
Boltzmann frequencies and phenotypic frequencies: (left) the average Boltzmann frequency *p*_*q*_ of an RNA secondary structure *q* over a sample of random sequences is plotted against the phenotypic frequency *f*_*q*_ of this structure, i.e. the probability that a random sequence folds into *q* as its minimum-free-energy structure. (right) same for the HP protein model. The same structures are shown as in Fig. 2C&F.

We can apply the insight from Fig. 3 to neutral-set-averages as follows: the sequences in a large neutral set are (almost) as diverse as arbitrary sequences, especially for RNA (SI section S5). Therefore, it is likely that the neutral-set-averages of Boltzmann frequencies in large neutral sets are similar to the averages over arbitrary sequences. This would mean that for structures *p* with high phenotypic frequency, we have *p*_*qp*_ *∝ f*_*q*_. Thus, we recover the existing result [1] that for an initial structure with a high neutral set, there is a high correlation between *ϕ* _*qp*_ and the phenotypic frequency of the new structure *f*_*q*_.

## III. CONCLUSIONS AND DISCUSSION

In this paper, we have shown that averages over Boltzmann ensembles capture two key GP map quantities: first, structures that have high Boltzmann probabilities in arbitrary sequences are more likely to fold and also have high phenotypic frequencies *f*_*q*_, in agreement with previous data for the RNAshapes model [6]. Secondly, if sequences folding into an initial structure *p* have a structure *q* as a high-Boltzmann-frequency alternative, then there are many mutations that transform *p* into *q*, in agreement with our data for the RNAshapes model [21]. This means that the existing principle of plastogenetic congruence [17] can be generalised from individual sequences to averages over the neutral set of structures, even though the mutational neighbourhood [1, 26] and the ensemble of suboptimal structures with high Boltzmann frequencies [20] differs markedly from sequence to sequence in a neutral set.

Intuitively, plastogenetic congruence only applies for well-behaved energy-functions, as in the theoretical calculations in ref [19], where free energies vary by a small amount after a single mutation. However, it should not apply when the energies of different structures exhibit jumps after single mutations. The HP model has such a well-behaved energy function because each residue has up to three contacts and so the free energy of a given structure can only change by up to three units after a mutation. For RNA structures, however, some mutations will cause free energy jumps in some structures by preventing a base pair from forming and thus making it impossible for the structure to fold at all. Despite this potential caveat, existing work on plastogenetic congruence [17] in RNA, as well as the data in this paper, demonstrates that the concept still holds for RNA. In fact, one classic paper [17] uses base pairing constraints to argue why rare mutational transitions might also be rare plastic transitions. Our results show that the concept holds for both the HP model and the RNA model, indicating that the free-energy jumps caused by base pairing constraints neither prevent nor enable plastogenetic congruence and its generalisations. The fact that plastogenetic congruence and similar effects are found in a range of systems [27–30], indicates that our results may also hold more broadly for a range of GP maps. In fact, one analysis of protein superfamilies even indicates that such concepts may apply when we consider different sequences from the same fold [31]. However, many of these analyses focus on continuous phenotypic and structural changes, whereas here we have focused specifically on the probabilities of obtaining specific discrete structural changes because these probabilities have been shown to be highly biased [12] and impact evolutionary processes [3]. Future work should investigate connections between these different perspectives on mutational changes.

One implication of our results is that structures that have high average Boltzmann probabilities and are thus likely to evolve as suboptimal structures, also have high *ϕ* _*qp*_ values and are thus more likely to evolve as mfe structures, as conjectured in Ancel and Fontana’s original paper on plastogenetic congruence [17]. This applies for arbitrary sequences, where the relevant quantities are *f*_*q*_ and *p*_*q*_, and for a given initial structure, when the relevant quantities are *ϕ* _*qp*_ and *p*_*qp*_. This finding is highly relevant in cases where the fitness of a sequence is not determined simply by the identity of its mfe structure, but instead depends on the Boltzmann frequency of the correctly folded structure, as was found in a large-scale experimental study on tRNAs [32]. In this case, the likelihood of evolving a given structure *q* would not be given by *ϕ* _*qp*_ and *f*_*q*_ values, but by a quantity that reflects the possibility that *q* could emerge as a suboptimal structure, i.e. a quantity related to our Boltzmann averages *p*_*q*_ or *p*_*qp*_. The close correspondence between our Boltzmann averages and existing GP map quantities means that previous work [4, 7], which has relied on mfe structures to predict which structures are likely to evolve, will continue to apply, even if *p*_*q*_ or *p*_*qp*_ are relevant, as in the computational models in ref [33], rather than the mfe-based quantities *f*_*q*_ or *ϕ* _*qp*_.

Our results have a practical application in sampling methods: usually, GP map analyses are restricted to short sequences because of the computational cost required to sample a sufficient number of sequences to estimate quantities like phenotypic frequencies and mutation probabilities *ϕ* _*qp*_. While specialised sampling algorithms have been developed to approximate phenotypic frequencies from particularly small samples for the RNA model [34, 35], Boltzmann averages could be a second option for approximating trends in *f*_*q*_ data and *ϕ* _*qp*_ data from a small sample for a broad range of GP maps. In this approach, more information is extracted from each sequence in the sample than if we limit ourselves to the mfe structure of each sequence: the additional information from the Boltzmann ensemble of folding could lower the well-known [36] sampling errors for low *f*_*q*_ and *ϕ* _*qp*_ values. This is especially useful in cases, where entire Boltzmann ensembles can be predicted as quickly as the mfe structure, for example in the HP model, where the mfe structure is usually identified by computing the energy of all folds (for example [10, 37]). In addition, the results may be useful when making inferences from partial experimental data on mutational changes and fluctuations.

## IV. METHODS

### A. RNA secondary structures

We use the ViennaRNA [9, 16, 38] package (version 2.4.14) with no isolated base pairs for structure predictions. We only consider sequences with unique mfe structures as folding (due to the discrete nature of the energy model, this is not guaranteed for all sequences [39]), but we do include sequences that have a structure without base pairs as their unique minimum-free-energy state. Due to the high number of 4^30^ ∝ 10^18^ sequences of length *L* = 30, we rely on sampling approaches.

*Sampling approaches used for the RNA data in Fig. 2*: *ϕ* _*pq*_ and *p*_*pq*_ for a given initial structure *p* are approximated from two separate sequence samples, which are both generated with our adaptation [22] of Weiß and Ahnert’s site-scanning method [35] (parameters: 100 site-scanning processing with 10^5^ steps and subsequent subsampling of one in 50 sequences). Our adaptation includes base pair swaps and is therefore more suitable for sampling from neutral sets, not just their connected components. This is repeated for a total of 50 initial structures *p*, which are randomly drawn out of all structures obtained from folding 10^8^ sequences - the random draw is weighted such that there are a similar number of structures with *x* stacks. Since the number of stacks is closely linked to neutral set sizes [7], this procedure ensures that a range of neutral set sizes are represented. We compute Boltzmann ensembles up to 15 kcal/mol above the mfe structure. To reduce sampling errors, we only plot values of *ϕ* _*pq*_ *≥*5*/*(3*nL*), *f*_*q*_ *≥*5*/*10^8^ and *p*_*pq*_ *≥*0.2*/n*, where *n* is the number of sequences in the sample and *L* the sequence length. Phenotypic frequencies are estimated by folding 10^8^ random sequences since this allows us to obtain both a full list of high-frequency structures and frequency estimates for these structures. To reduce sampling errors, we only plot values of ≥5*/*10^8^. Our sampling methods are tested in section S2 of the SI.

*Sampling approaches used for the RNA data in Fig. 3*: the phenotypic frequencies of the 50 structures *p* from above are estimated using the approach of Jörg et al. [34], but with isolated base pairs switched off. This method does not test for uniqueness of mfe structures, but the impact of this is minimal for neutral set sizes [40]. The Boltzmann averages *p*_*f*_ are based on 10^7^ random sequences.

### B. HP protein model

Our data relies on a full enumeration of all HP sequences of length *L* = 25 and their folded structures on a compact lattice, using a simple energy model [41] with a contact energy of one unit for two hydrophobic residues and no energy contribution otherwise. We follow the steps outlined by Greenbury et al. [1, 37, 42] in the construction of the GP map and in the convention of disregarding sequences with multiple mfe structures, and test our methods against their data [42]. However, in this paper we treat two structures as distinct structures if they have reversed directionality, i.e. if the structure looks identical except with the N-terminus and C-terminus swapped, as in ref [43]. This ensures consistency with the RNA folding model, where information on directionality in the folded structure is also retained. *f*_*q*_ and *ϕ* _*qp*_ values are determined exactly from this data. *p*_*qp*_ values are approximated by the average over 10^3^ sequences drawn with replacement from the neutral set of *p*. Boltzmann-averages over arbitrary sequences are approximated by the average over 10^5^ sequences. We set the reduced temperature to *k*_*B*_*T* = 0.5 (relative to the HP interaction strength) since this represents a realistic middle ground between *k*_*B*_*T* = 0.1 on the one extreme, where the protein has no plasticity and typically spends *>* 99% of time in the ground state, and *k*_*B*_*T* = 1 on the other extreme, where the ground state accounts for less than 5% of the Boltzmann ensemble of a typical sequence. However, our results also hold for *k*_*B*_*T* = 0.1 and *k*_*B*_*T* = 1 (shown in section S6 of the SI).

## V. ACKNOWLEDGEMENTS

The authors would like to thank J. Blundell and T McLeish for discussion. N. Martin acknowledges funding from the Gates Cambridge Trust, the Winton Programme for the Physics of Sustainability and the Issachar Fund. S.E.A. was supported by the Gatsby Charitable Foundation with grant no. PTAG/021. (2015).

## Supplementary Information

### S1 Boltzmann frequencies and phenotypic frequencies - without mfe structures

In the main text, we found that the average Boltzmann probability of a structure *q* over a set of random sequences is correlated with the phenotypic frequency *f*_*q*_ of this structure. One trivial explanation for this finding could be the following: if the Boltzmann distribution of each sequence was dominated by the mfe structure, then the average Boltzmann probability would simply be a reflection of how often structure *q* is the mfe structure, which is exactly the definition of the phenotypic frequency *f*_*q*_. To see if this trivial relationship is the reason for our observed correlation, we repeated an adjusted version of the analysis without this potential confounding factor (Fig. S1): we computed the average Boltzmann probability of a structure *q* over a set of random sequences but excluded sequences where it was the mfe structure or one of several degenerate mfe structures. We still find a correlation between the average Boltzmann probability of *q* and its phenotypic frequency for both RNA structures and the HP protein model, indicating that our observed correlation is not just due to the dominance of mfe structure in the Boltzmann ensemble.

**Figure S1:**
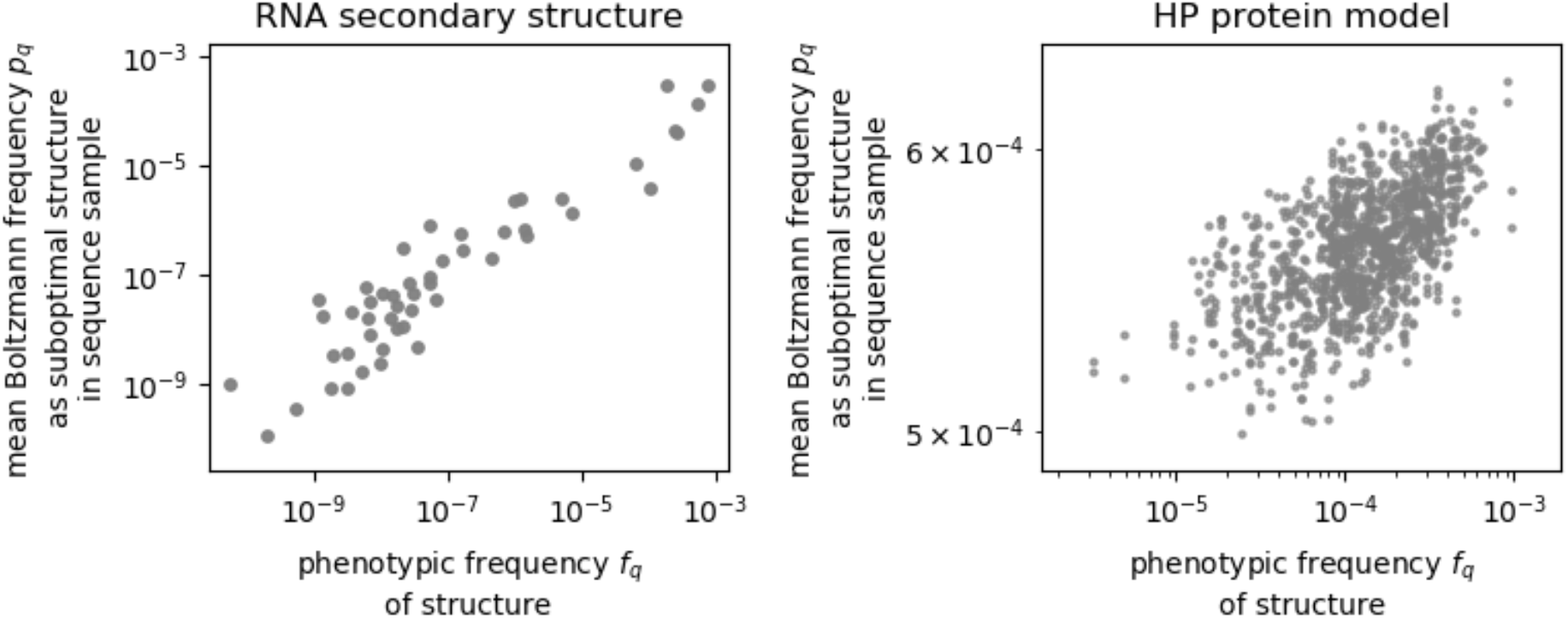
Boltzmann frequencies and phenotypic frequencies: (left) the average Boltzmann frequency of each RNA secondary structure q is estimated from a sample of random sequences - here a sequence is only included in the calculation if q is not the mfe structure or one of several degenerate mfe structures for this sequence. This quantity is plotted against the phenotypic frequency f_q_ of this structure, i.e. the probability that a random sequence folds into q as the unique mfe structure. (right) same for the HP protein model.

### S2 Sampling methods

Our *ϕ* _*qp*_ and *p*_*qp*_ data for RNA secondary structures is based on a sequence sample of 2 × 10^5^ sequences per initial structure *q*. This sample was generated using an adaptation of Weiß and Ahnert’s [1] site-scanning method, as described in the main text. To ensure that this method introduced no systematic biases, we adapt an analysis in the SI of our previous paper [2]: we obtain a second sequence sample for each neutral set and compare the *ϕ* _*qp*_ and *p*_*qp*_ values from both approaches. As a second method (‘sequence sampling’ approach), we simply generate random sequences and keep them if they happen to fold into any of our initial structures *p*. If, after sampling 10^9^ random sequences, we have at least 10^4^ sequences for a given initial structure *p*, we take the first 10^4^ sequences for *p* and compute *ϕ* _*qp*_ and *p*_*qp*_ for that sequence sample. With these parameters, the ‘sequence sampling’ approach will generate a sufficient number of sequences for structures *p* with high phenotypic frequency of approximately *f*_*p*_ ⪆ 10^4^*/*10^9^ = 10^5^. This data is shown in Figs S2 & S3. We find that the data shows excellent agreement at high values of *ϕ* _*qp*_ and *p*_*qp*_ and some sampling errors for lower values. Thus, up to expected sampling errors that play a role when *ϕ* _*qp*_ ⪅ 10^4^ and *p*_*qp*_ ⪅ 10^4^, the ‘sequence sampling’ approach, which samples each sequence with the same probability and has therefore no systematic bias, gives the same *ϕ* _*qp*_ and *p*_*qp*_ values as we computed from the modified site-scanning method. The modified site-scanning method is used throughout the paper because it works for any structure *p*, and not just for those with high phenotypic frequency *f*_*p*_.

**Figure S2:**
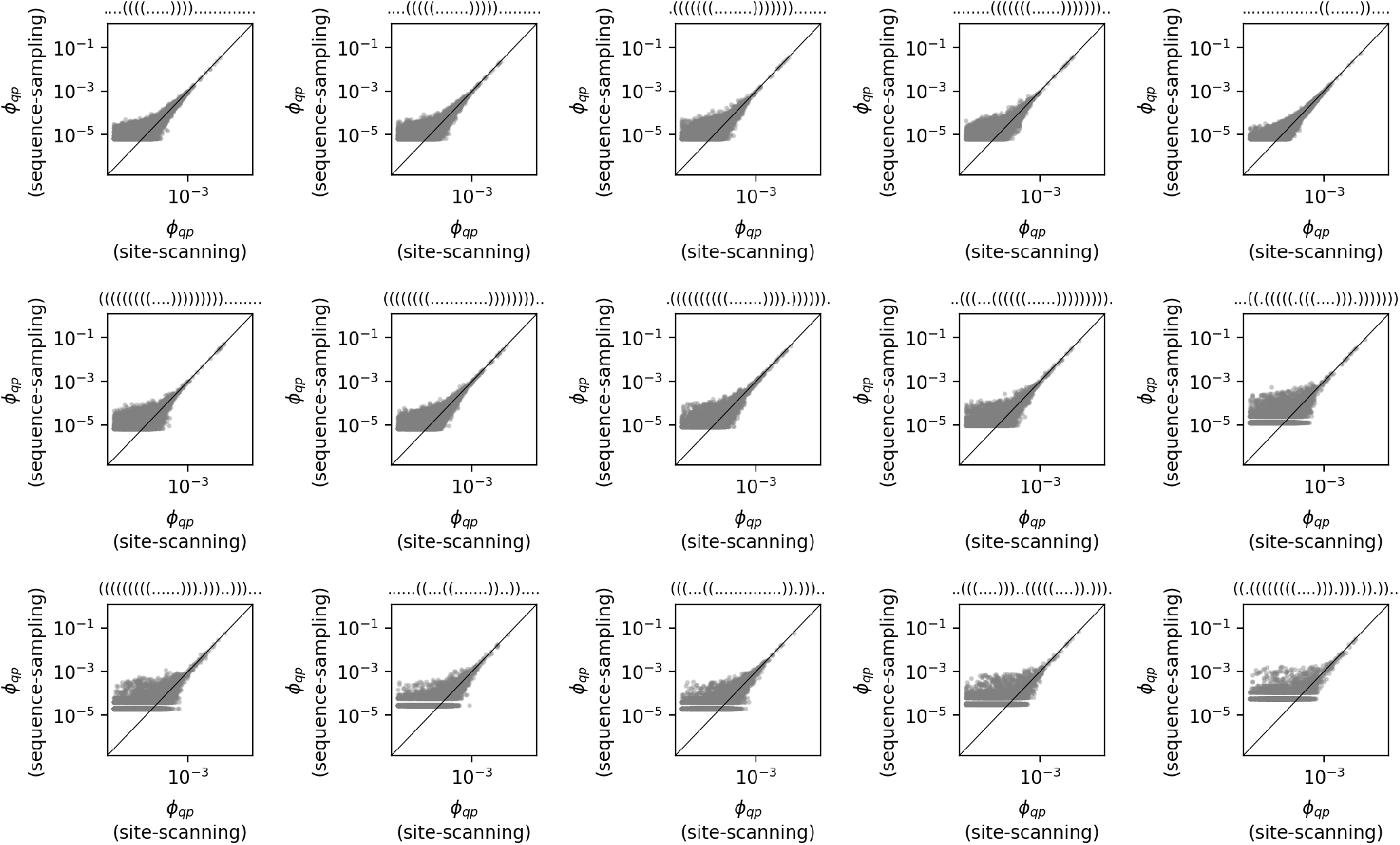
Comparison of the ϕ _qp_ data for RNA from two sampling approaches: in order to test our sample-based ϕ _qp_ data, we compute ϕ _qp_ from two different sequence samples based on two different approaches (an adaptation of Weiß and Ahnert’s [1] site-scanning method and a simple ‘sequence sampling’ approach, as described in the text). Each subplot shows the ϕ _qp_ data for a given initial structure p, which is given in the subplot title. The ‘sequence sampling’ approach can only be applied if p has a high phenotypic frequency f_p_, and therefore only a subset of the 50 structures p used in the main text are shown. For these structures, the ϕ _qp_ values from the two methods are in good agreement, except for sampling errors for low ϕ _qp_ values of ϕ _qp_ ⪅ 10^4^, which are consistent with the finite sample size of 10^4^ in the sequence-sampling approach.

**Figure S3:**
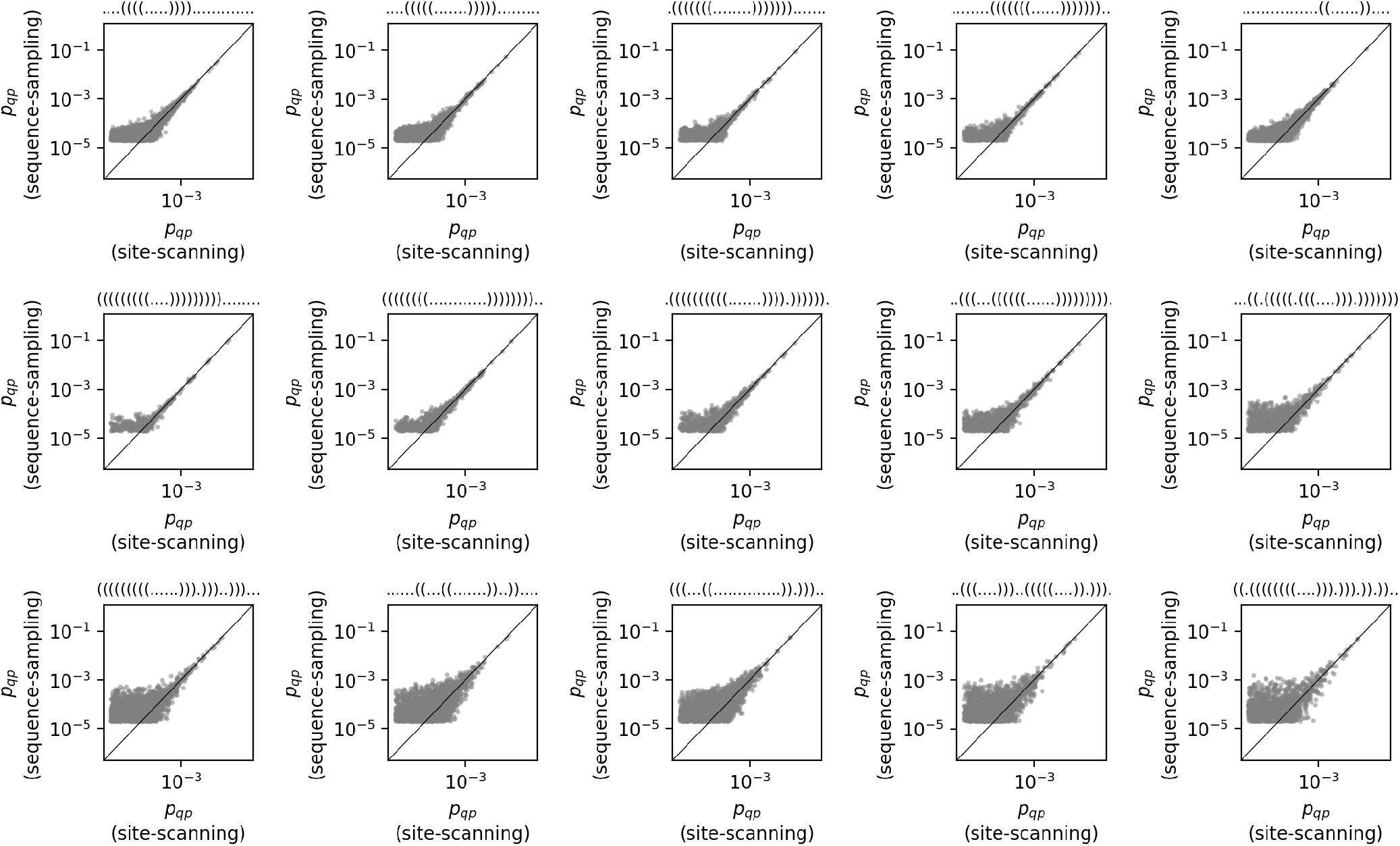
Comparison of the p_qp_ data for RNA from two sampling approaches: same as Fig. S2 but for p_qp_ instead of ϕ _qp_.

### S3 *ϕ* _*qp*_ and Boltzmann frequencies - further examples

In the main text, we only had space to show the data for a single initial structure *p* and included summary data for further choices of *p*. Here, we show the full data for further choices of the initial structure *p*: first, we plot the phenotypic frequency of the new structure *f*_*q*_ against the phenotypic mutation probability *ϕ* _*qp*_ (i.e. the approach from ref [3, 4]) for 25 RNA (Fig. S4) and 35 protein structures (Fig. S6). Secondly, we present the corresponding data based on our new approach, i.e. using the average Boltzmann frequency of the new structure in the neutral set of the initial structure *p*_*qp*_ as in ref [2], in Figs S5 & S7. These plots confirm the conclusions from the main text: our new approach is a much better indicator of *p*_*qp*_ differences, both for proteins and RNA.

**Figure S4:**
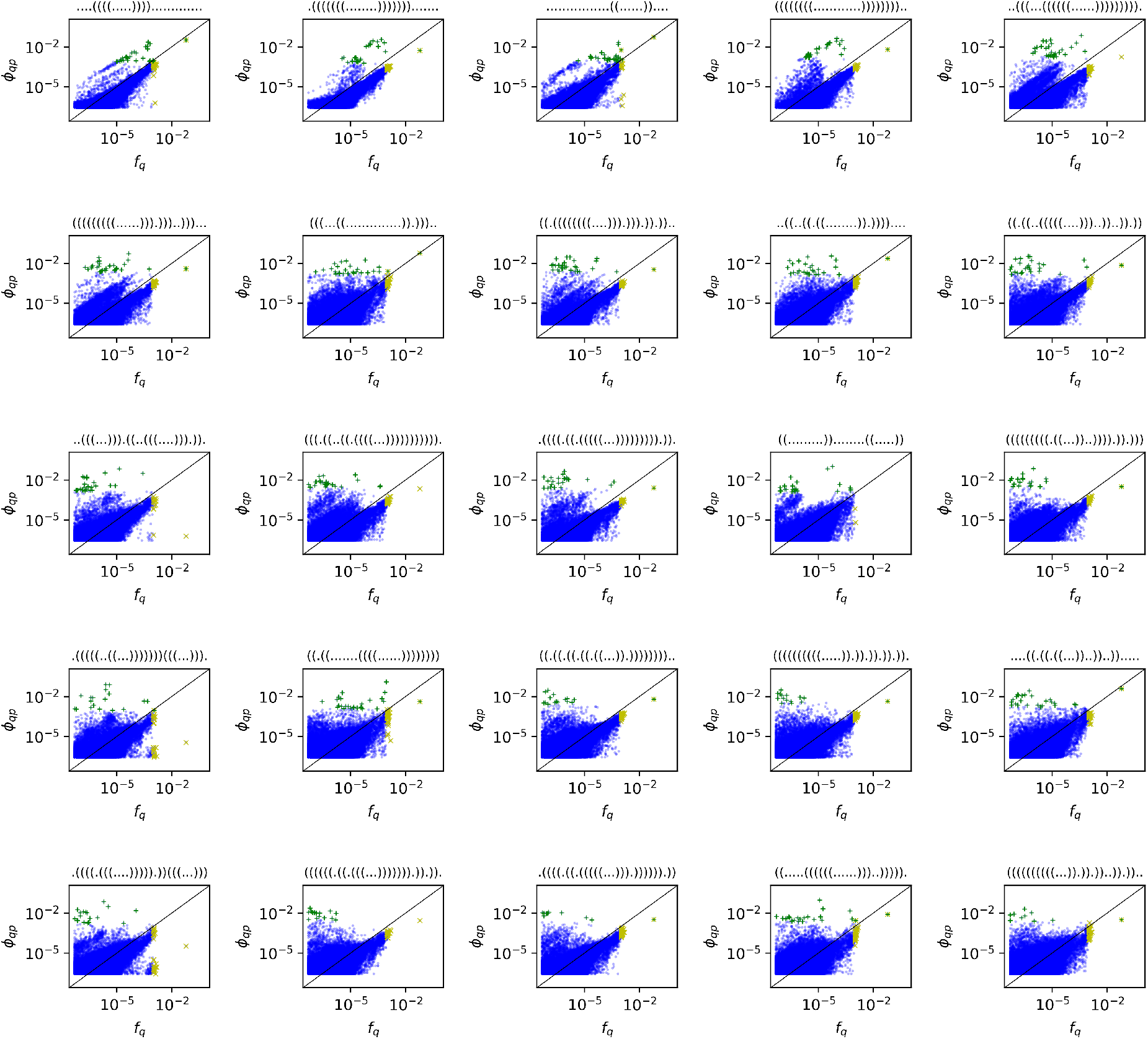
Existing approach for 25 different initial RNA structures. p: each plot uses a different initial structure p. The ϕ _qp_ values for mutating from this initial structure p to a new structure q are plotted against the phenotypic frequency f_q_ of the new structure q. In order to visualise, which data points correspond to the top-30 highest values, the top-30 highest ϕ _qp_ values are indicated by green ‘+’ scatter points and the top-30 highest f_q_ values are indicated by yellow ‘x’ scatter points. We see that there is little overlap between these two groups and so the structures with the top-30 highest f_q_ values are not the same structures as those with the top-30 highest ϕ _qp_ values, consistent with the statistics in Fig. 2C in the main text. Every second structure from the sorted list of 50 RNA structures from Fig. 2C in the main text is shown and each structure p is given in the plot title in dot-bracket notation.

**Figure S5:**
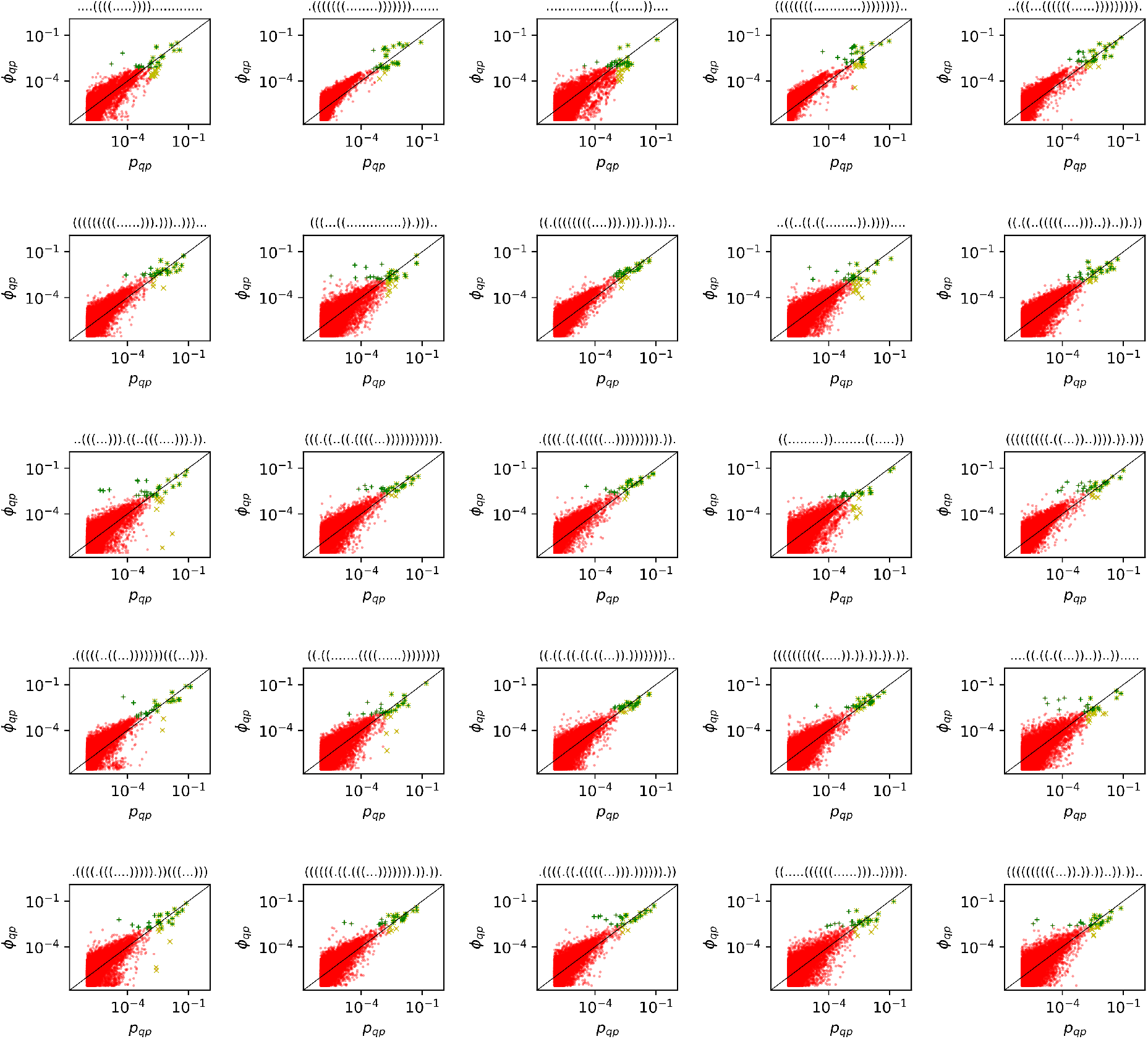
New approach for 25 different initial RNA structures p: each plot uses a different initial structure p. The ϕ _qp_ values for mutating from this initial structure p to a new structure q are plotted against the Boltzmann frequency p_qp_ of the new structure q averaged over sequences in the neutral set of the initial structure p. In order to visualise, which data points correspond to the top-30 highest values, the top-30 highest ϕ _qp_ values are indicated by green ‘+’ scatter points and the top-30 highest p_qp_ values are indicated by yellow ‘x’ scatter points. We see that there is considerable overlap between these two groups and so many of the structures with the top-30 highest p_qp_ values are among the top-30 highest ϕ _qp_ values, consistent with the statistics in Fig. 2C in the main text. Every second structure from the sorted list of 50 RNA structures from Fig. 2C in the main text is shown and each structure p is given in the plot title in dot-bracket notation.

**Figure S6:**
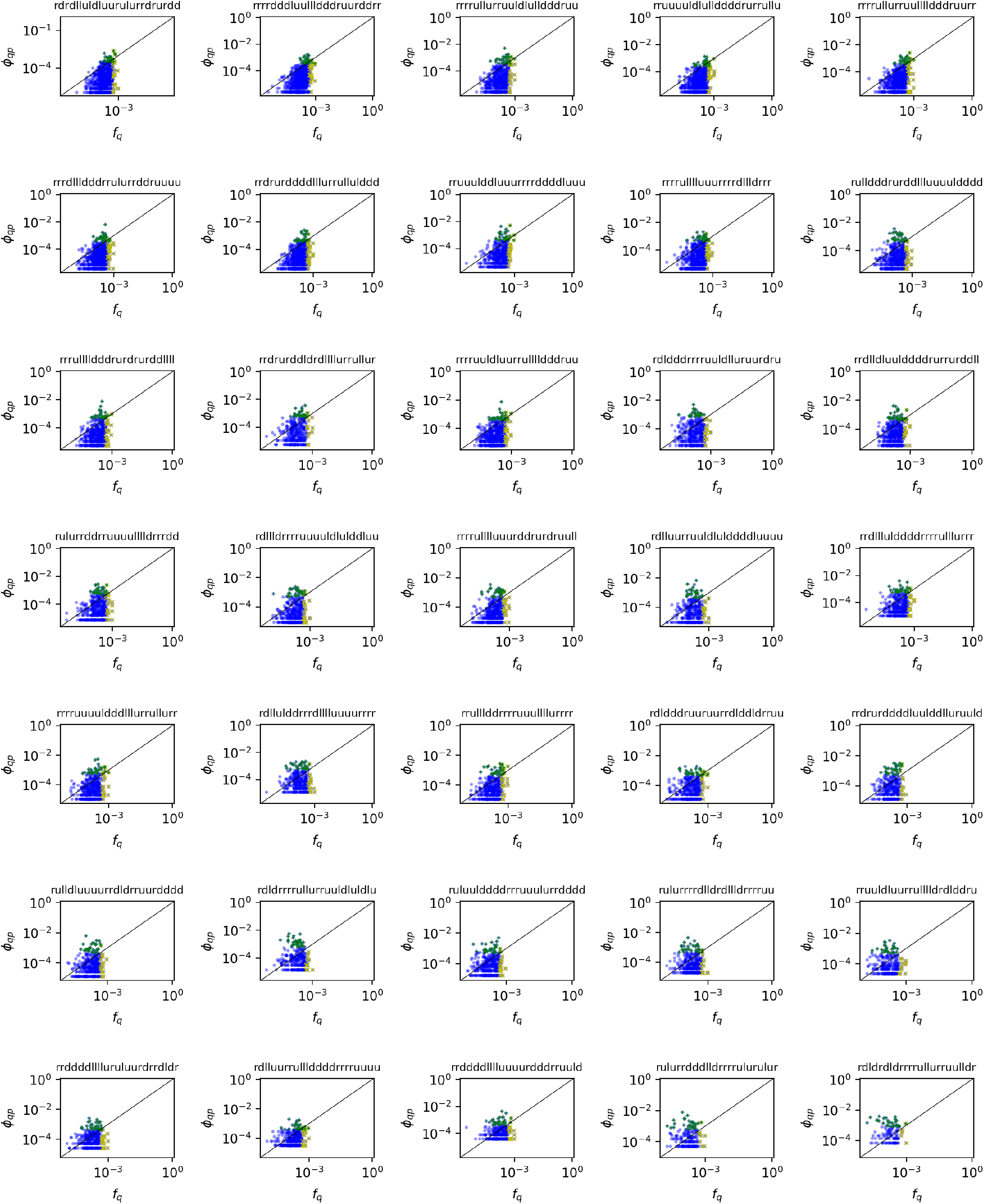
Existing approach for 35 different initial HP protein structures p: each plot uses a different initial structure p. The ϕ _qp_ values for mutating from this initial structure p to a new structure q are plotted against the phenotypic frequency f_q_ of the new structure q. The 35 structures are chosen at regular intervals from the sorted list of 1081 HP structures from Fig. 2F in the main text and each structure p is given in the plot title (the notation follows the chain along the lattice and ‘r’, ‘l’, ‘u’ and ‘d’ stand for a move ‘right’, ‘left’, ‘up’ and ‘down’).

**Figure S7:**
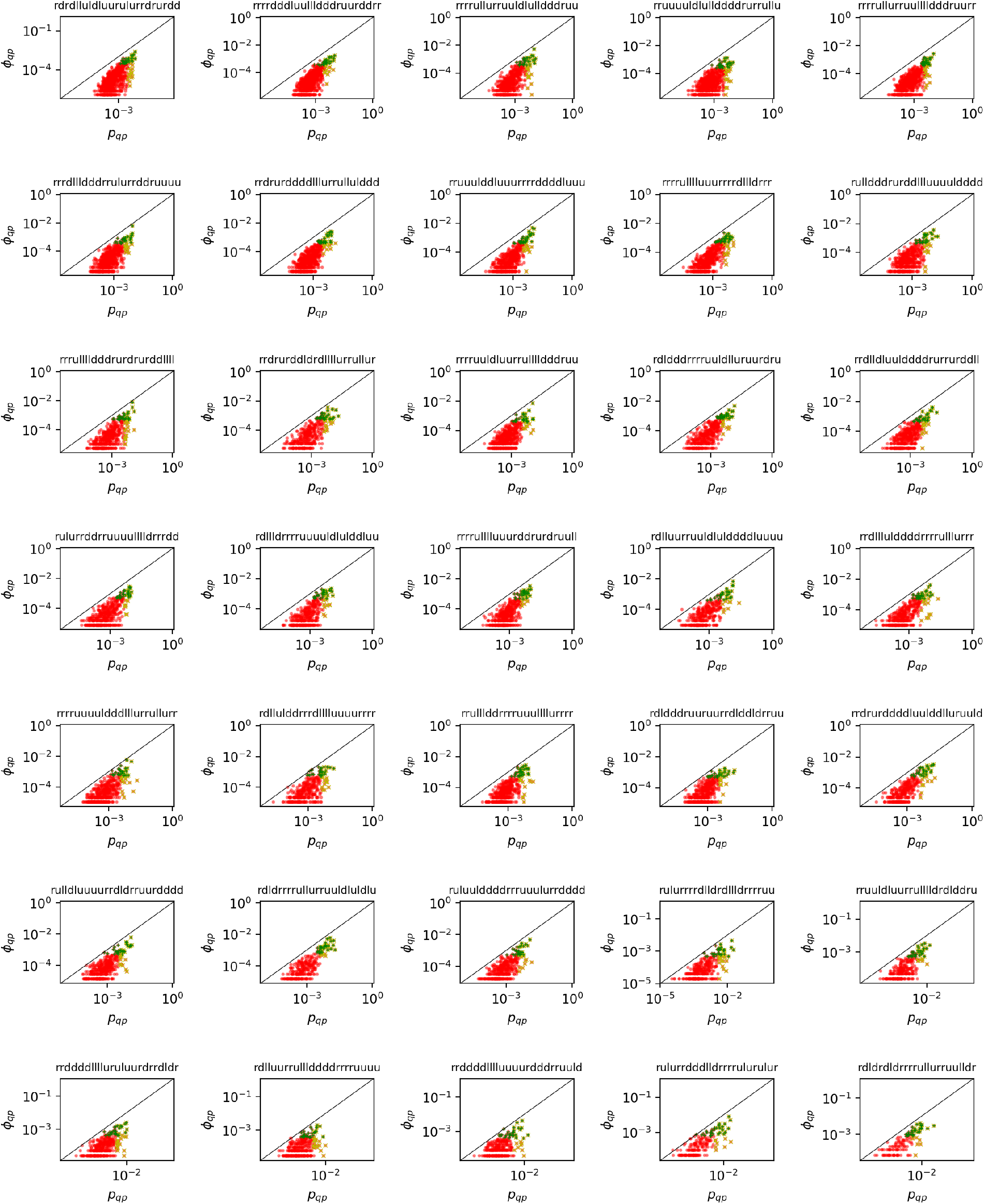
New approach for 35 different initial HP protein structures p: each plot uses a different initial structure p. The ϕ _qp_ values for mutating from this initial structure p to a new structure q are plotted against the Boltzmann frequency p_qp_ of the new structure q averaged over sequences in the neutral set of the initial structure p. The 35 structures are chosen at regular intervals from the sorted list of 1081 HP structures from Fig. 2F in the main text and each structure p is given in the plot title (the notation follows the chain’s walk along the lattice and denotes each step by ‘r’, ‘l’, ‘u’ or ‘d’, which stand for ‘right’, ‘left’, ‘up’ and ‘down’).

### S4 Comparison with further alternative approaches

#### S4.1 *ϕ* _*qp*_ and structural similarity between *p* and *q*

**Figure S8:**
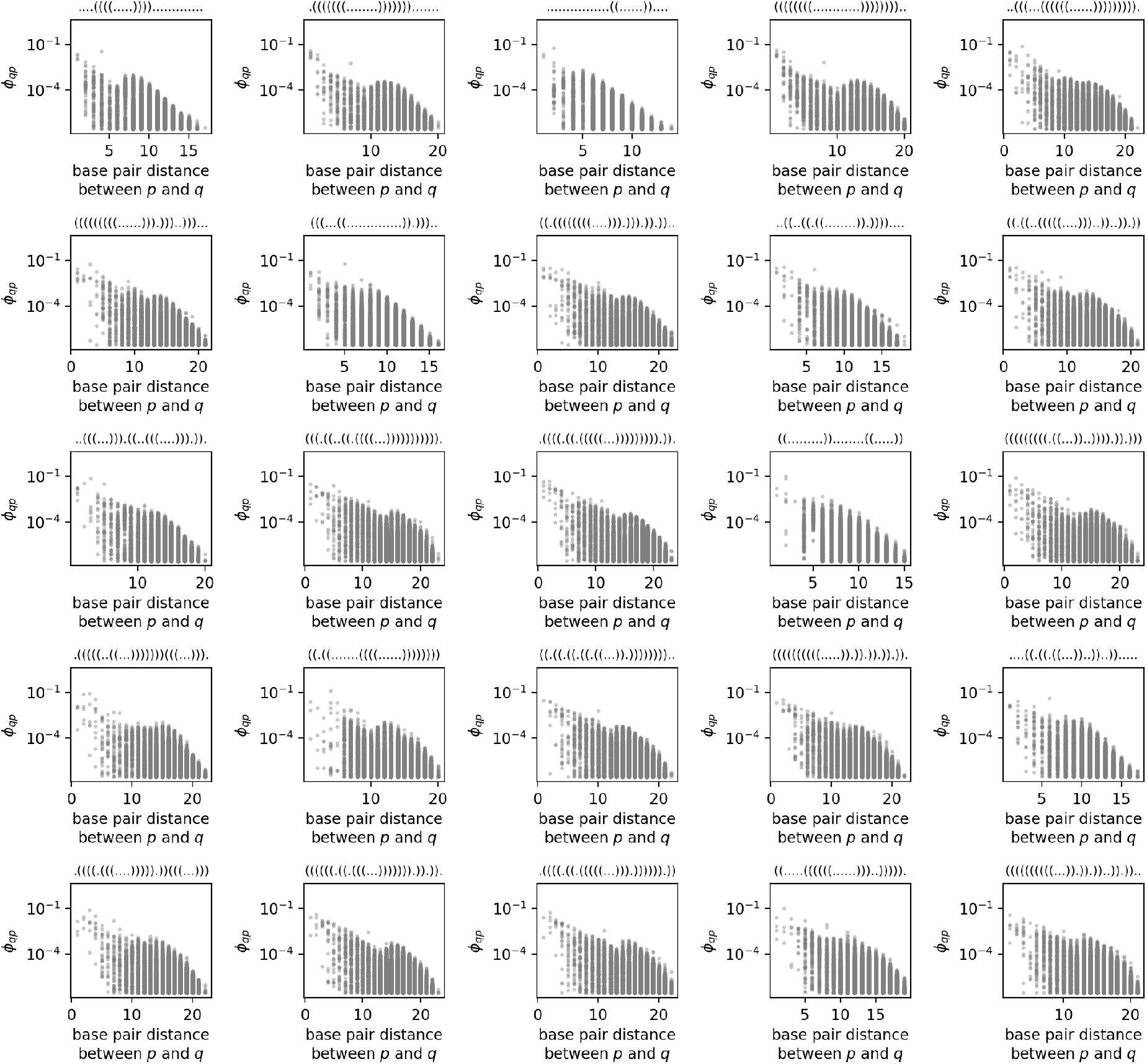
Mutational transition probability ϕ _qp_ versus structural difference between RNA structures p and q: as in the previous figures, each subplot focuses on one specific initial structure p that is given in the subplot title. The ϕ _qp_ values for structural changes to a new structure q are plotted against the base pair distance between p and q, which is calculated using the ViennaRNA package [5]. We find that likely phenotypic changes (i.e. high ϕ _qp_ values) tend to correspond to small structural distances, but not vice versa: for a given base pair distance, we often find a range of ϕ _qp_ values, including likely and unlikely ones. Note that because of the logarithmic scale and sampling method, there may be further structural changes with zero or low ϕ _qp_ values.

One alternative approach for approximating the likelihood of specific structural changes is to postulate that small structural changes are more likely to occur than large changes and therefore *ϕ* _*qp*_ values should be linked to the structural similarity between *p* and *q* (for example ref [4, 6]).

We test this hypothesis in Figs S8 - S11, using the same structures as examples as in Figs S4 - S7, so that the plots can be compared directly. In Fig. S8, *ϕ* _*qp*_ values in the RNA model are plotted against the base pair distance between *p* and *q*, an established structural distance definition implemented in the ViennaRNA package [7]. Similarly, in Fig. S9, *ϕ* _*qp*_ values in the HP protein model are plotted against the contact map overlap between *p* and *q*, an established structural similarity metric for proteins [8]. Fig. S10 repeats the analysis based on a metric that counts whether the sequence positions that are in the core of structure *p* are also located in the core of the new structure *q*. This metric was chosen because previous work on slightly different HP models had shown that the identity of core and surface sites is a structural quantity that is especially important for the GP map [9]. Finally, we also computed for RNA, what fraction of sequences in the neutral set of the initial structure *p* are compatible with the new structure *q* (Fig. S11). In this context ‘compatible’ means that a sequence can form the base pairs required for folding into *q* [10], but might do so with only an infinitesimal Boltzmann frequency. We also plot a closely related quantity in Fig. S12: what fraction of sequences that is compatible with *p* is also compatible with *q*. This final measure was suggested in ref [10], who in turn cite J. Weber’s 1997 PhD thesis. We find that none of these similarity measures or distance measures is a good predictor of *ϕ* _*qp*_: they all capture some *ϕ* _*qp*_ differences, and the base pair distance correctly identifies some highly likely transitions, but in all cases, there is a range of *ϕ* _*qp*_ for a given structural similarity value. This is exactly what we would expect from Fontana and Schuster’s classification of likely and unlikely mutational changes [11] in their framework, adding or deleting a single base pair at the end of a stack is considered a likely structural change, but other additions or deletions of single base pairs do not appear among the different types of likely structural changes [11]. Similarly, deletions of stacks are considered a likely change, but additions of stacks are not [11], even though both are the same in terms of the base pair distance between the old and the new structure. Thus, out of all possible structural transitions with a base pair distance of *x*, only some are likely to occur, and we would expect a range of *ϕ* _*qp*_ values at a constant structural similarity, which is exactly what we observe.

For symmetric similarity measures like the base pair distance, there is another argument that explains why *ϕ* _*qp*_ values cannot be approximated well: the base pair distance is the same, regardless of whether we compare *p* to *q* or *q* to *p*, but *ϕ* _*qp*_ and *ϕ* _*pq*_ can differ by several orders of magnitude if the neutral set sizes of *p* and *q* differ [10].

One way of understanding, why our Boltzmann-based approach is better at approximating *ϕ* _*qp*_ trends goes as follows: likely structural transitions tend to be to structures that are structurally similar to the initial structure, but not *all* structures that are structurally similar have high transition probabilities. Similarly, Boltzmann states close to the mfe structure tend to be structurally similar to the mfe structure [12], but presumably not all Boltzmann states that are structurally similar to the mfe structure are among the energetically lowest-lying states.

Finally, the fact that the shared number of compatible sequences only gives an upper bound for *ϕ* _*qp*_ values is consistent with our previous result for neutral set sizes, where the number of base pairs and hence compatible sequences was found to be only weakly predictive of neutral set size [13]. We found that thermodynamic considerations are also necessary since the fraction of compatible sequences that actually adopt the structure as a mfe structure differs considerably from structure to structure [13].

**Figure S9:**
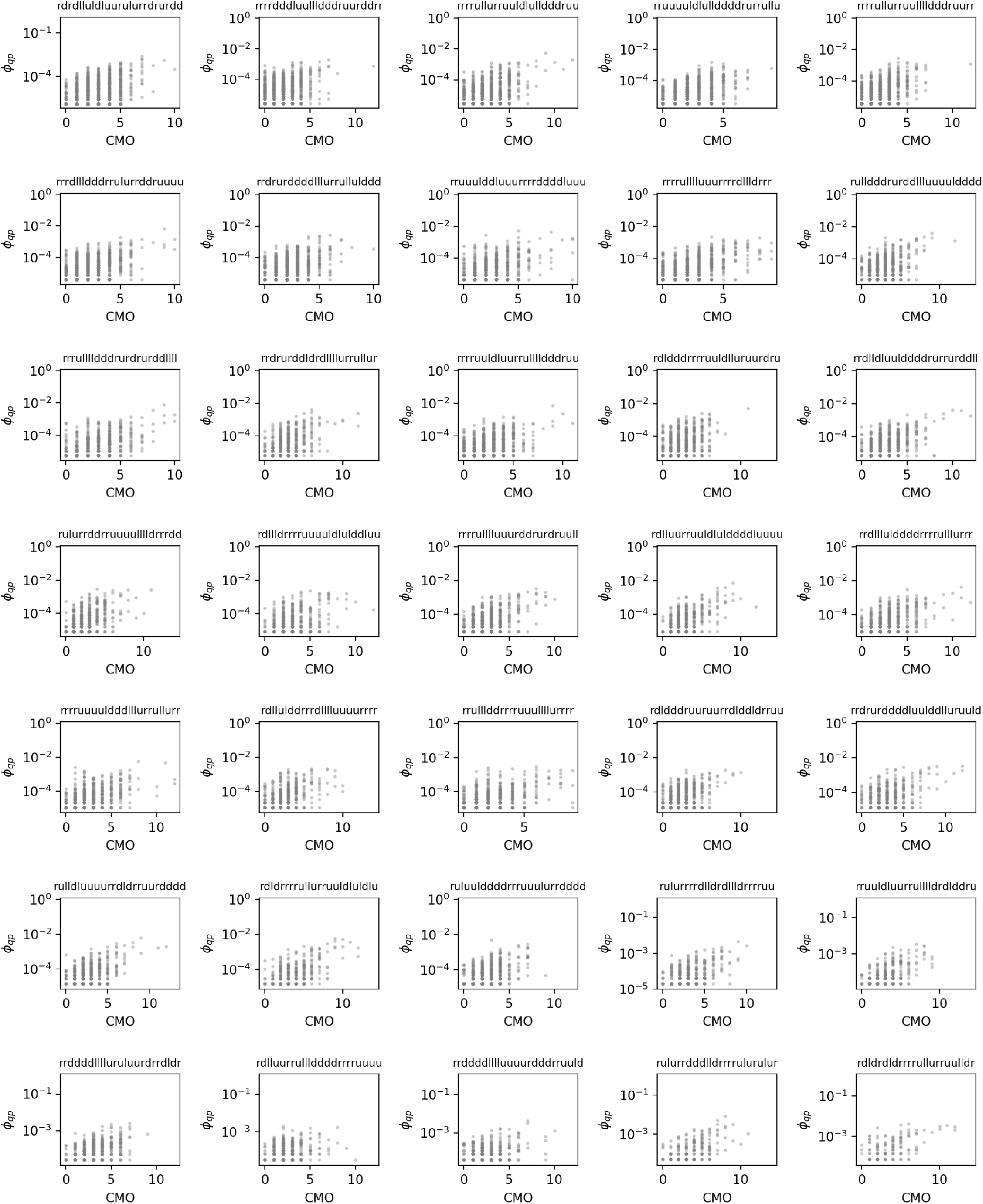
Mutational transition probability ϕ _qp_ versus structural difference between HP protein structures p and q: as in the previous figures, each subplot focuses on one specific initial structure p that is given in the subplot title. The ϕ _qp_ values for structural changes to a new structure q are plotted against the contact map overlap (CMO) between p and q. We find only a weak trend.

**Figure S10:**
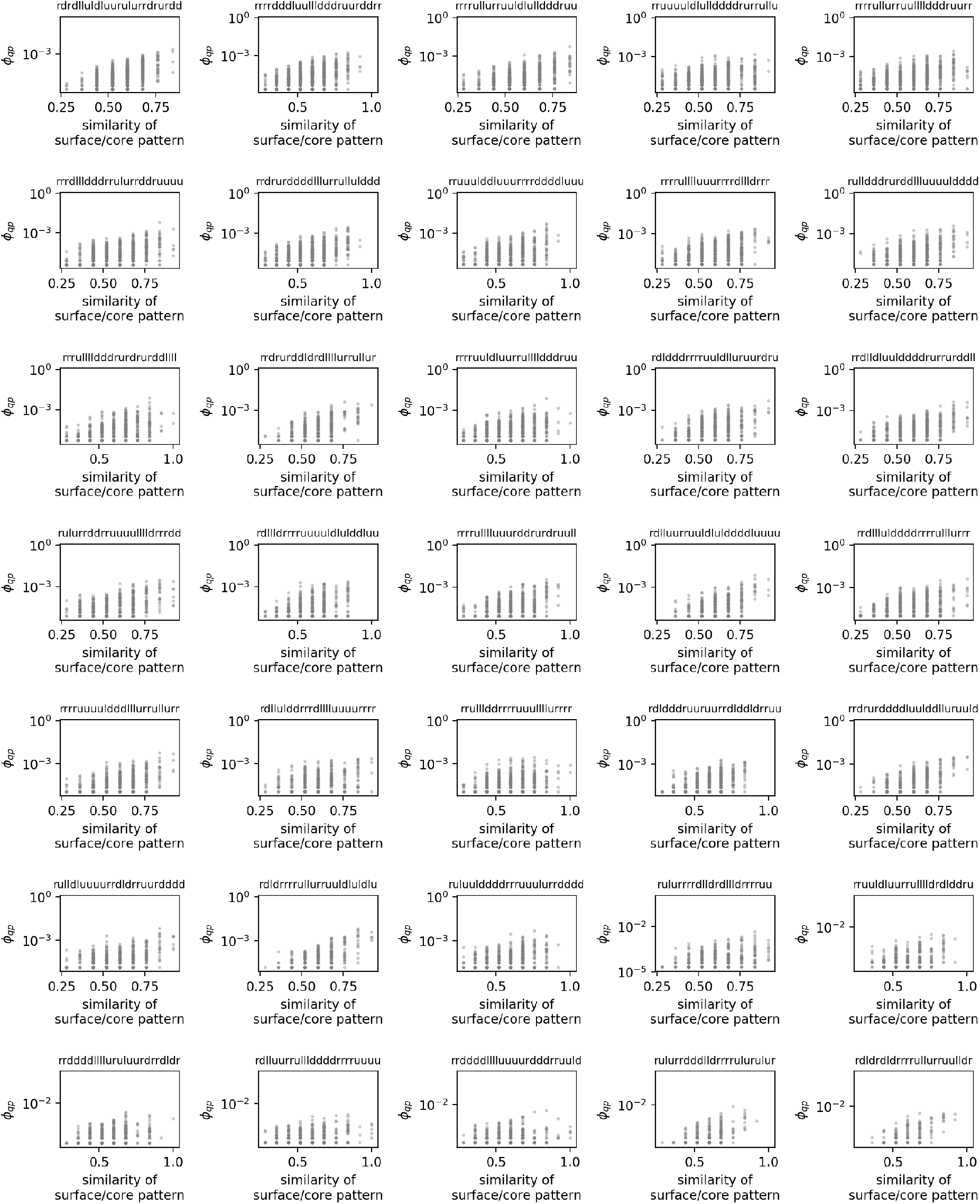
Mutational transition probability ϕ _qp_ versus structural distance between HP protein structures p and q: as in the previous figures, each subplot focuses on one specific initial structure p that is given in the subplot title. The ϕ _qp_ values for structural changes to a new structure q are plotted against the fraction of sites that share the same structural context in both structures (i.e. they are either in the core of both structures p and q or on the surface of both structures).

**Figure S11:**
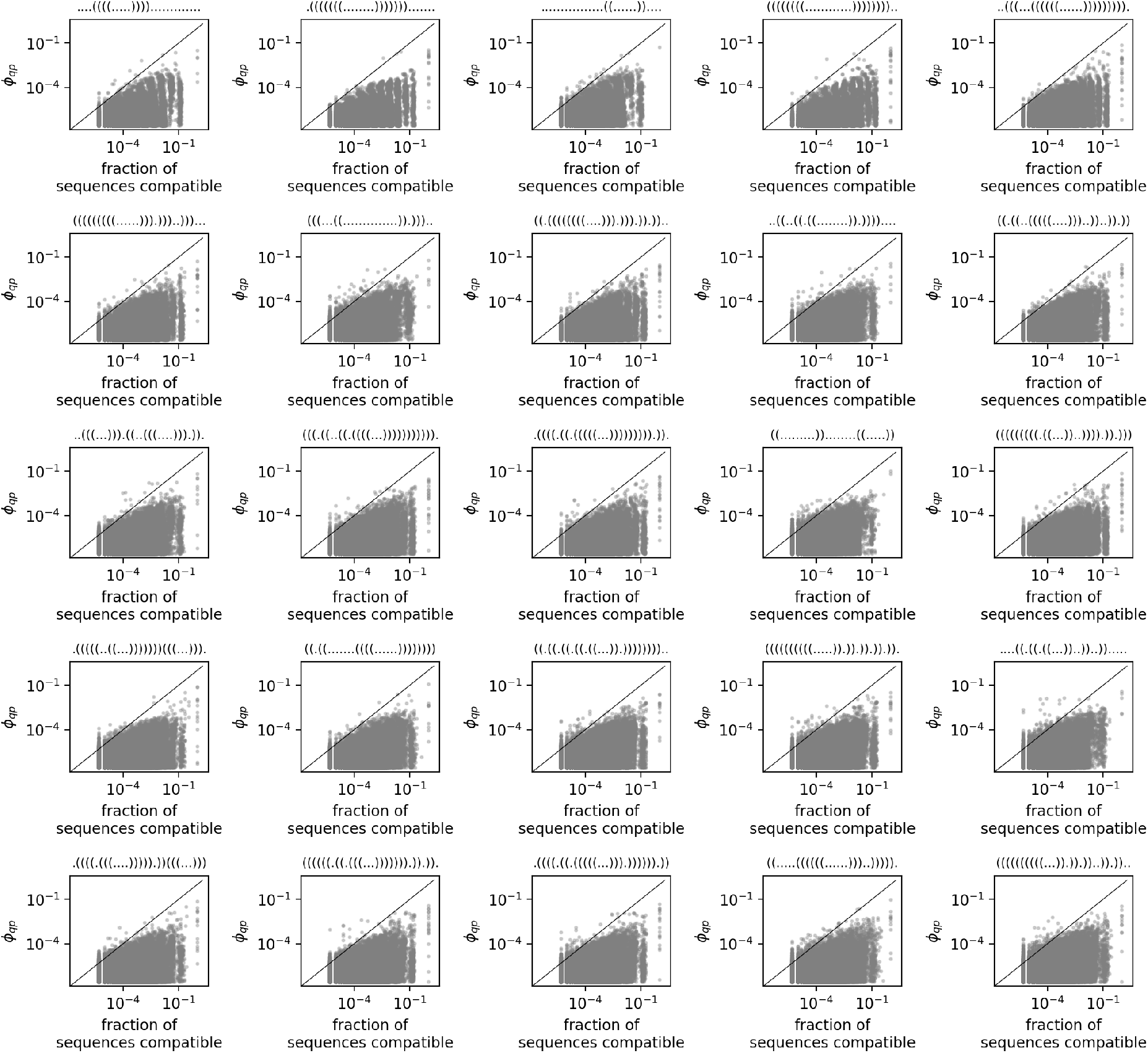
Mutational transition probability ϕ _qp_ versus overlap in the number of compatible sequences between RNA structures p and q: as in the previous figures, each subplot focuses on one specific initial structure p that is given in the subplot title. The ϕ _qp_ values for structural changes to a new structure q are plotted on the y-axis. The data on the x-axis quantifies, what fraction of sequences in the neutral set of p are compatible with q. We find that likely phenotypic changes (i.e. high ϕ _qp_ values) tend to correspond to high overlaps, but not vice versa: for a given overlap, we find a range of ϕ _qp_ values, including likely and unlikely ones.

**Figure S12:**
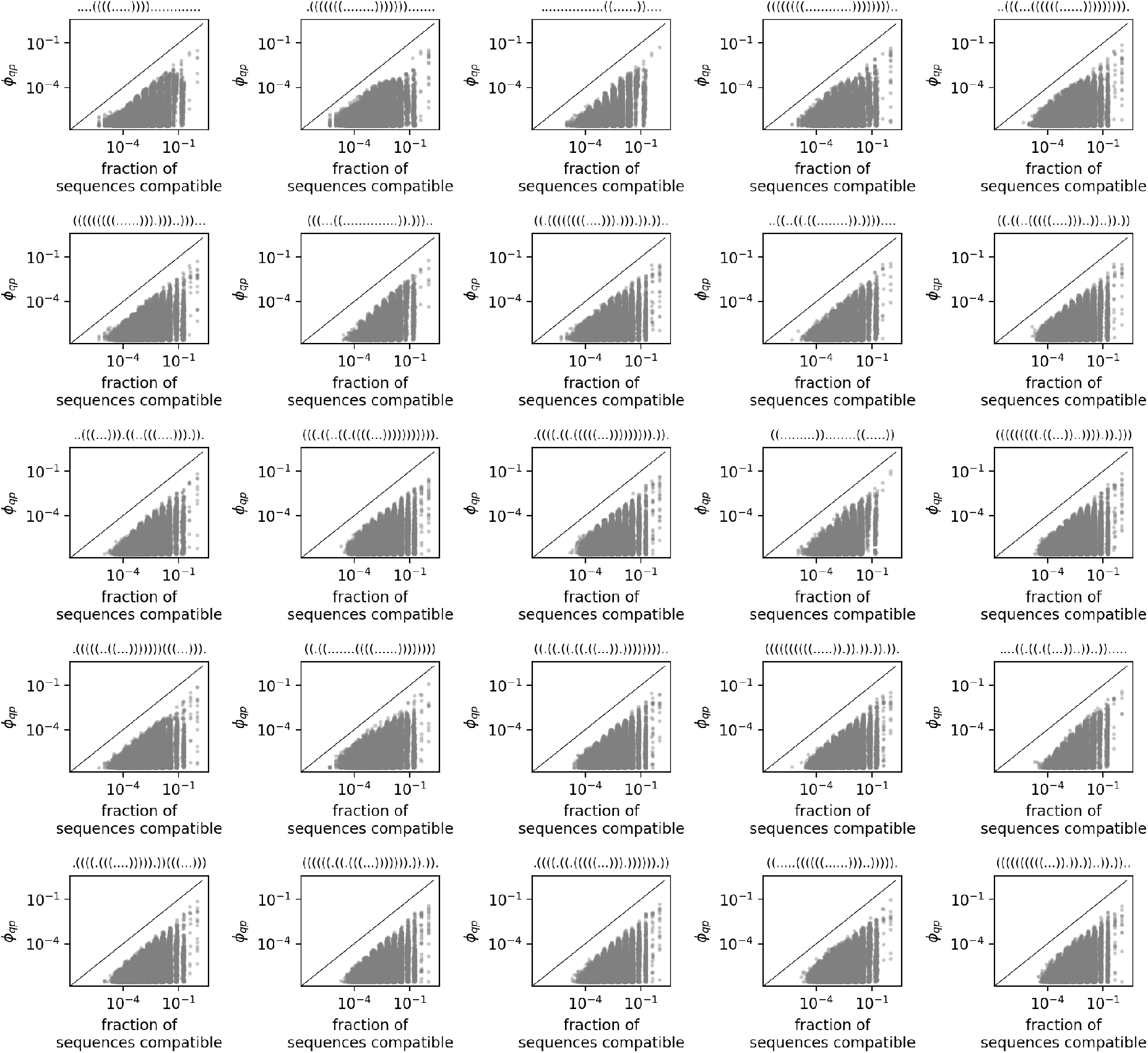
Mutational transition probability ϕ _qp_ versus overlap in the number of compatible sequences between RNA structures p and q: as in the previous figures, each subplot focuses on one specific initial structure p that is given in the subplot title. The ϕ _qp_ values for structural changes to a new structure q are plotted on the y-axis. The data on the x-axis quantifies, what fraction of sequences that are compatible with p are also compatible with q (i.e. similar to Fig. S11, but evaluated relative to all compatible sequences of p and not only the neutral set of p). We find that likely phenotypic changes (i.e. high ϕ _qp_ values) tend to correspond to high overlaps, but not vice versa: for a given overlap, we find a range of ϕ _qp_ values, including likely and unlikely ones.

#### S4.2 Information-theoretic approach

We also apply the recent approach [14] based on algorithmic information theory (AIT), again applying it to the same structures and *ϕ* _*qp*_ values as those in Figs S4 - S7. The AIT approach postulates that an upper bound to the log of the *ϕ* _*qp*_ values can be approximated by the conditional complexity of *q* given *p*, a measure of how much information is needed to describe *q* if *p* is known [14]. We computed this conditional complexity using the same compression-based approach as the authors (from ref [15]) and show this data for RNA in Fig. S13 and for the HP model in Fig. S14. We find that the relationship between *ϕ* _*qp*_ values and conditional complexities has a log-linear upper bound for RNA, as postulated in ref [14]. However, because this is only an upper bound, there are many phenotypes, for which we would predict high *ϕ* _*qp*_ values based on the conditional complexities, but which actually have low *ϕ* _*qp*_ values. Our new approach (Fig. S5) shows a much clearer correlation with *ϕ* _*qp*_ values. For the HP protein model, the AIT approach does not seem to give a clear trend at all (Fig. S14). However, it is possible that this is simply because there is no clear method of calculating conditional complexities for the HP models - here, we simply used the up-down notation as shown in the subplot titles as an input for the complexity calculation, and simply represent each of the four letters by a binary number (‘00’ for ‘right’, ‘01’ for ‘down’, ‘10’ for ‘left’, ‘11’ for ‘up’). It is possible that a different linear representation of HP structures exists that might give a clearer trend, but this has not been explored since the AIT approach has not been applied to lattice proteins yet, neither in the *ϕ* _*qp*_ paper [14] nor in a bigger analysis that applied a related AIT approach to phenotypic frequencies for a range of models [16].

#### S4.3 Systematic comparison

In sections S4.1 & S4.2, we have seen that similarity measures and AIT arguments capture *ϕ* _*qp*_ differences less clearly than our new approach. To make this comparison more quantitative, we repeat the analysis for further initial structures *p* and compare the methods directly: we apply this to all 1081 structures for the HP model and the same sample of 50 RNA structures as in the main text. The approach from the main text, where we evaluated, how many of the top-30 *ϕ* _*qp*_ values for this *p* can be identified with each approach, cannot be applied here for two reasons: first, the data may contain several identical values since the contact map overlap, conditional complexity and base pair distance are all discrete. In this case, we cannot clearly determine, which structures are predicted to have the 30 highest values because there may be ties. Secondly, while our sampling errors return all structures with high values of *ϕ* _*qp*_, *f*_*q*_ and *p*_*qp*_, we do not know, which of the ≈ 10^6^ possible RNA structures^1^ have the optimum base pair distance or conditional complexity without going through a full list of structures, which is infeasible.

**Figure S13:**
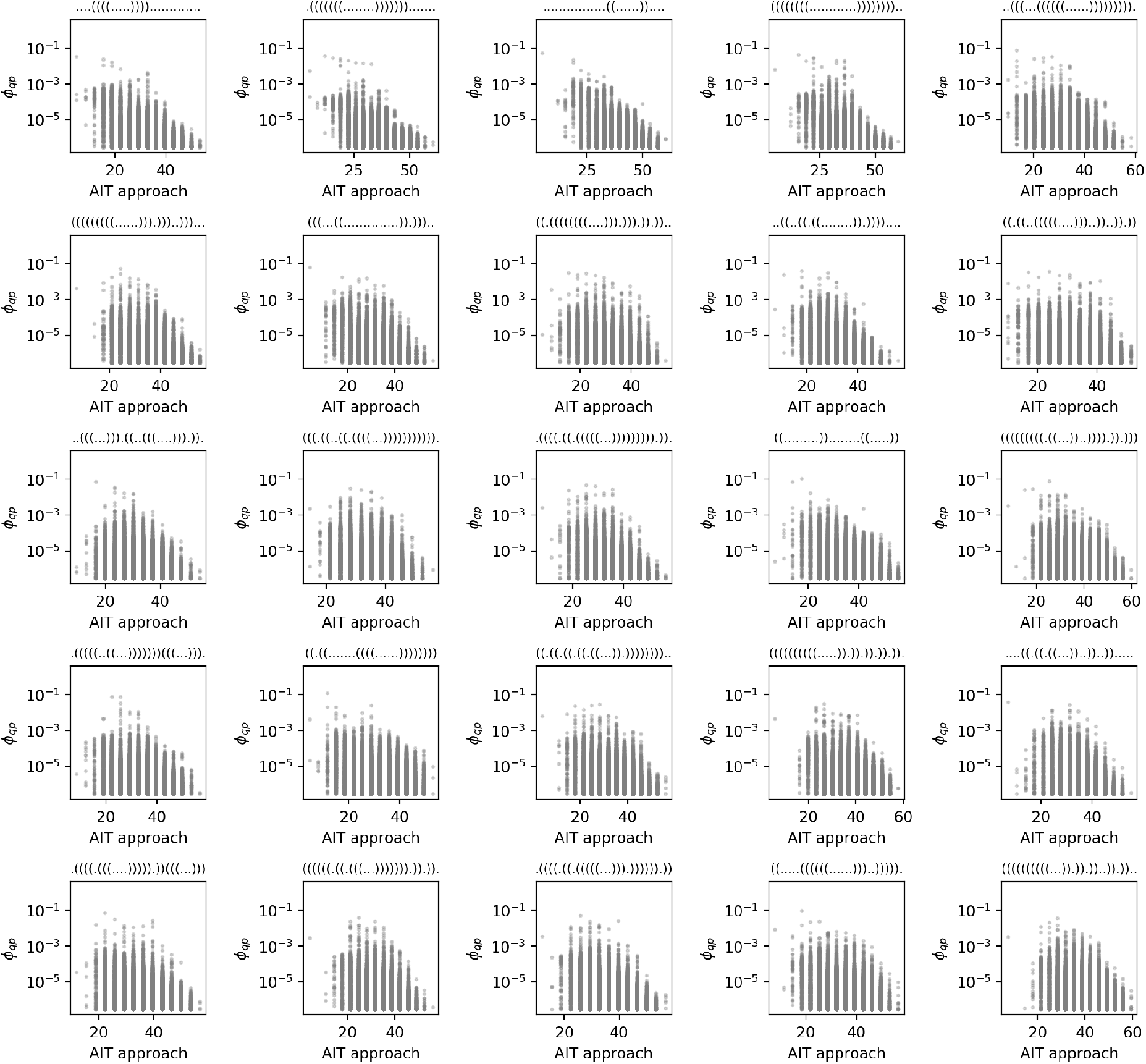
Mutational transition probability ϕ _qp_ versus conditional complexity of q given p for RNA structures: as in the previous figures, each subplot focuses on one specific initial structure p that is given in the subplot title. The ϕ _qp_ values for structural changes to a new structure q are plotted against the conditional complexity 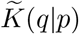 of q given p for RNA structures, as suggested in ref [14]. All complexity calculations follow the methods published by the same authors [15]. We find that likely phenotypic changes (i.e. high ϕ _qp_ values) tend to correspond to low conditional complexity values, but not vice versa: for a given conditional complexity value, we often find a range of ϕ _qp_ values, including likely and unlikely ones. This is consistent with the theory in ref [14], which predicts a log-linear upper bound for this data, but no lower bound.

**Figure S14:**
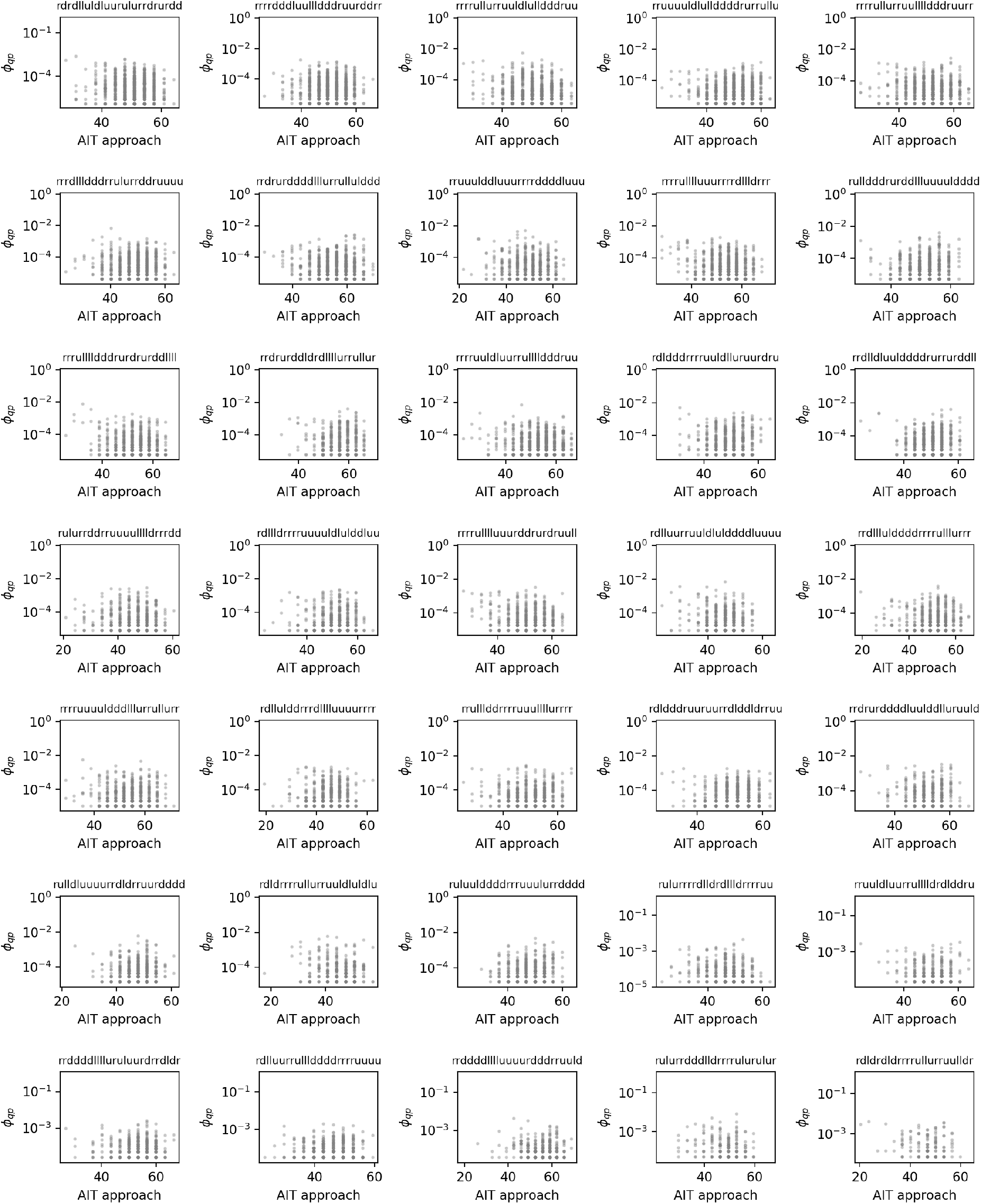
Mutational transition probability ϕ _qp_ versus conditional complexity of q given p for HP protein structures: same as Fig. S13, but for HP protein structures. Since there is no existing method of converting each HP structure into a binary string for the complexity calculations, we simply map the sequence of steps to the left/right/up/down that defines a HP structure onto a binary string, as described in the text. Here we do not see a clear log-linear upper bound, such as the one predicted in ref [14].

**Figure S15:**
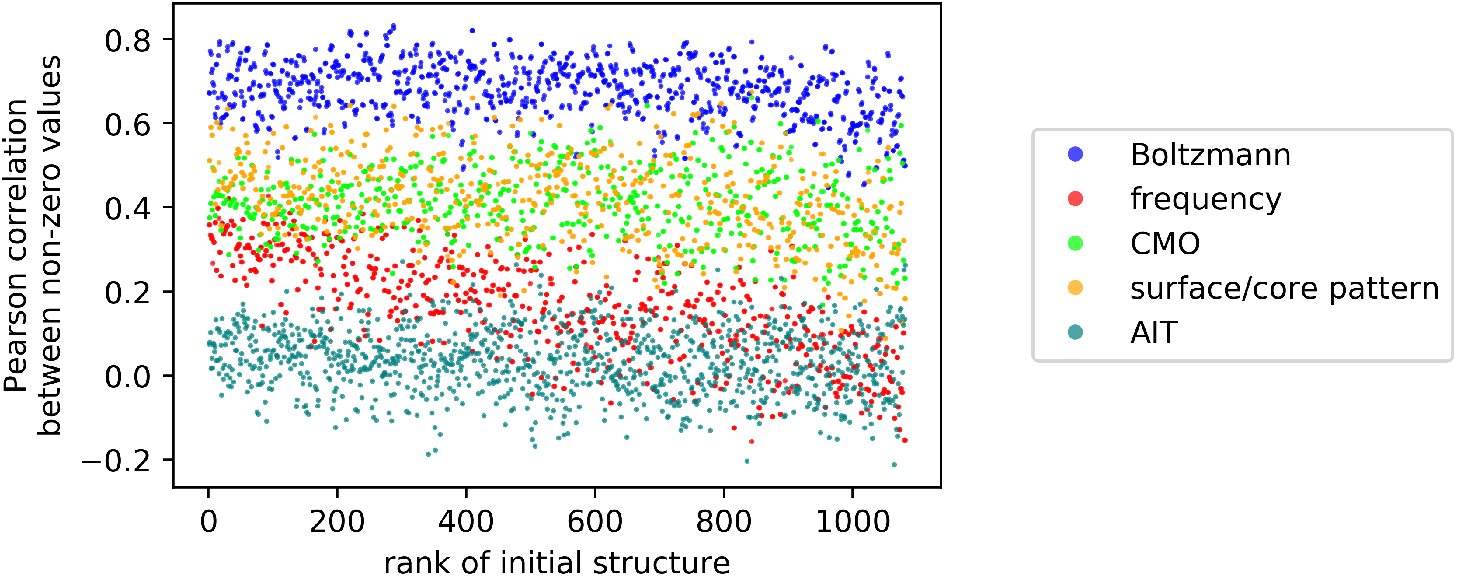
Comparison of all approaches for HP protein structures: for each of the 1081 initial structures p, each denoted by a unique rank as in the main text (x-axis), we compare the five different approaches by plotting the Pearson correlation coefficient between the data from each approach and the ϕ _qp_ values (y-axis). The five different approaches are: the new approach based on Boltzmann averages p_qp_ (blue), the existing approach [4] based on phenotypic frequencies f_q_ (red), the structural similarity based on contact map overlaps (lime), the structural similarity based on the similarity in the surface/core pattern (orange) and the AIT-based approach [14] (teal). The Pearson correlation coefficient is computed on the log-log scale (lin-log for the AIT and structural similarity calculations), and so phenotypes q with estimated ϕ _qp_ = 0 are not included. For methods which are expected to give a negative correlation (for example the AIT approach), we show the strength of the negative correlation.

**Figure S16:**
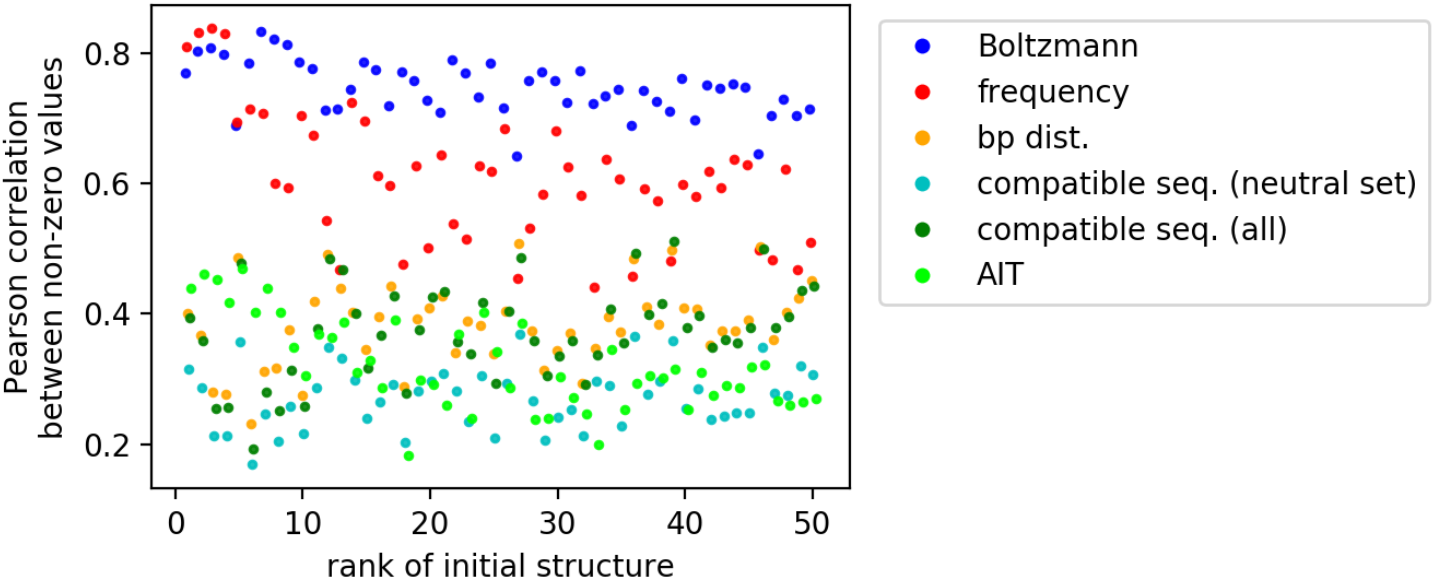
Comparison of all approaches for RNA secondary structures: same as Fig S15, but for the 50 initial RNA structures p from the main text. Here, the six different approaches are: the new approach based on Boltzmann averages p_qp_ (blue), the existing approach [4] based on phenotypic frequencies f_q_ (red), the structural similarity based on base pair distances (orange), the overlap in compatible sequences relative to the neutral set of p (cyan), the overlap in compatible sequences relative to the compatible sequences of p (green) and the AIT-based approach [14] (lime).

Therefore we choose a different way to quantify how well each approach captures *ϕ* _*qp*_ values: we compute the Pearson correlation coefficient between the predicted data and the *ϕ* _*qp*_ on a log-log scale (lin-log for the AIT approach, contact map overlap and base pair distance), to quantify the extent to which we can predict log *ϕ* _*qp*_ differences with each approach. The disadvantages of this method (and the reason why it is not used in the main text) is that because of the log-log scaling, zero values cannot be included and so cases, in which *ϕ* _*qp*_ = 0 and *f*_*q*_ *»* 0 are not accounted for. In addition, structures with low *ϕ* _*qp*_ values and therefore large sampling errors will lead to larger artefacts in the Pearson correlation coefficients than in the method used in the main text. However, we present the Pearson correlation data in spite of these caveats for two reasons: it provides an alternative to the method used in the main text and can be applied to data from all approaches.

Thus, we quantify the performance of each approach with Pearson correlation coefficients (Fig. S16 for RNA and Fig. S15 for the HP protein model): we find that, while the contact map overlap for HP proteins are a better indicator of likely phenotypic transitions (i.e. high *q>*_*qp*_ values) than phenotypic frequencies, our new approach gives the most accurate results. The only exception is RNA, where the existing approach based on phenotypic frequencies outperforms our new Boltzmann-frequency-based approach for a small number of high-frequency structures, consistent with ref [4]’s claim that this existing approach works best if the initial structure *p* has high phenotypic frequency.

### S5 Sequence diversity in neutral sets

**Figure S17:**
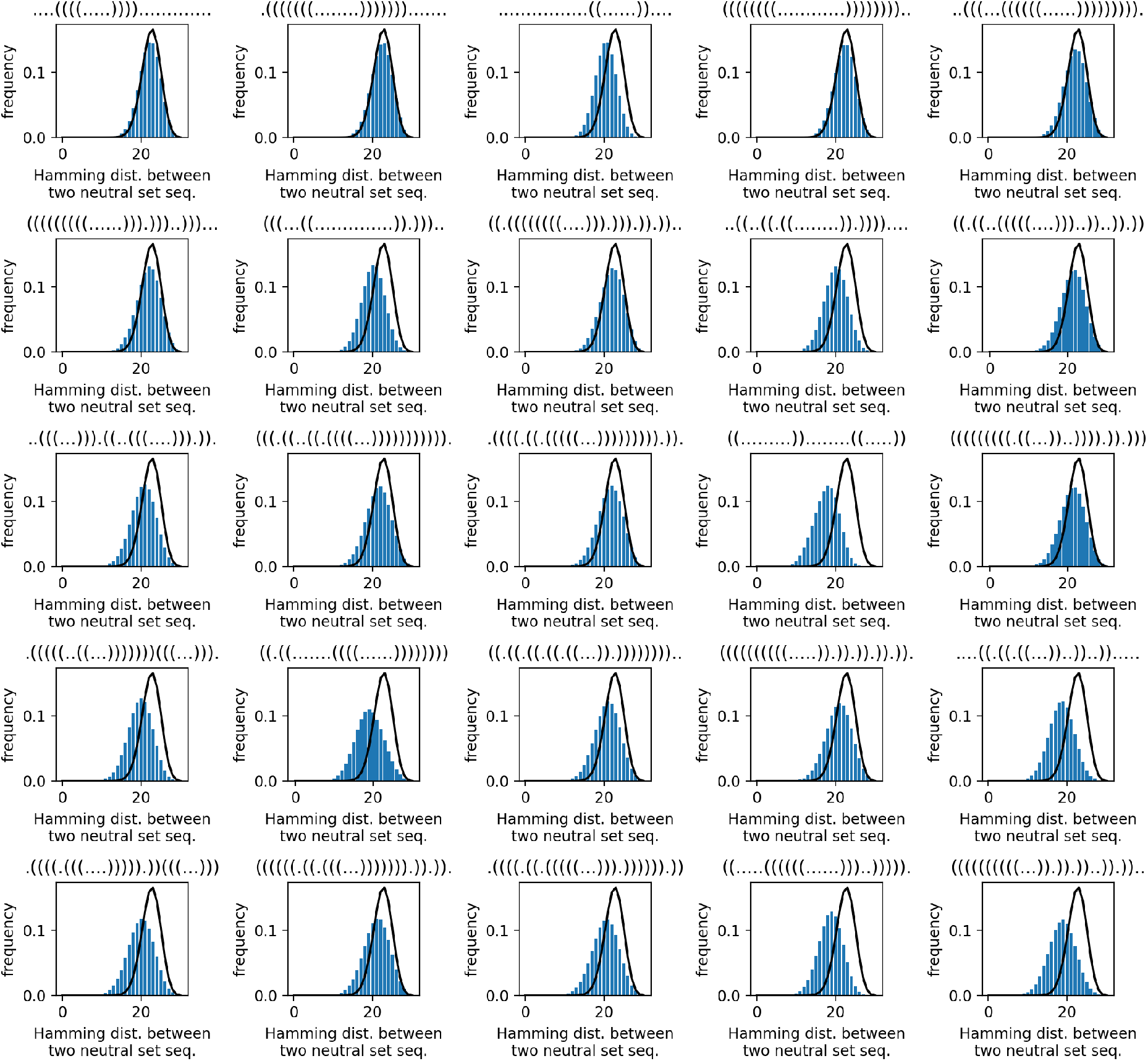
Diversity of sequences within several RNA neutral sets: each plot focuses on the neutral set a different initial structure p (using the same selection as the figures above). For each neutral set, we plot the Hamming distance distribution between different sequences in our neutral set (blue histogram), together with the Hamming distance distribution of random sequences (black line), as in our previous work on RNAshapes [2]. This data is based on 10^4^ sequence per neutral set taken from the sequence samples used for the p_qp_ calculations.

To get an understanding of how averages over random sequences (such as *p*_*q*_) relate to averages over neutral sets (such as *p*_*qp*_), we analyse, how the diversity of sequences in a neutral set differs from the diversity of random sequences. This analysis has already been performed for RNAshapes in our previous work [2], and further related quantities had been discussed in the past (for example [18–20]), but here we quantify this diversity systematically for the sequence lengths and models used in this analysis. For RNA (Fig. S17), we find that in large neutral sets, the distribution of sequence similarities is almost the same as the distribution among random sequences from different neutral sets, as previously found for the RNAshapes model [2]. In smaller neutral sets, sequences tend to have a higher similarity and thus lower Hamming distance. However, in all the neutral sets in Fig S17, the typical Hamming distance is still *>* 17, which is of the same order of magnitude as the Hamming distance between two random sequences (22.5). For HP proteins (Fig. S18), we find that two sequences from the same neutral set are more similar than two random sequences, but again there is some diversity within a neutral set: in the neutral sets shown here, two sequences typically have a Hamming distance between seven and eleven, compared to 12.5 between two random HP sequences.

**Figure S18:**
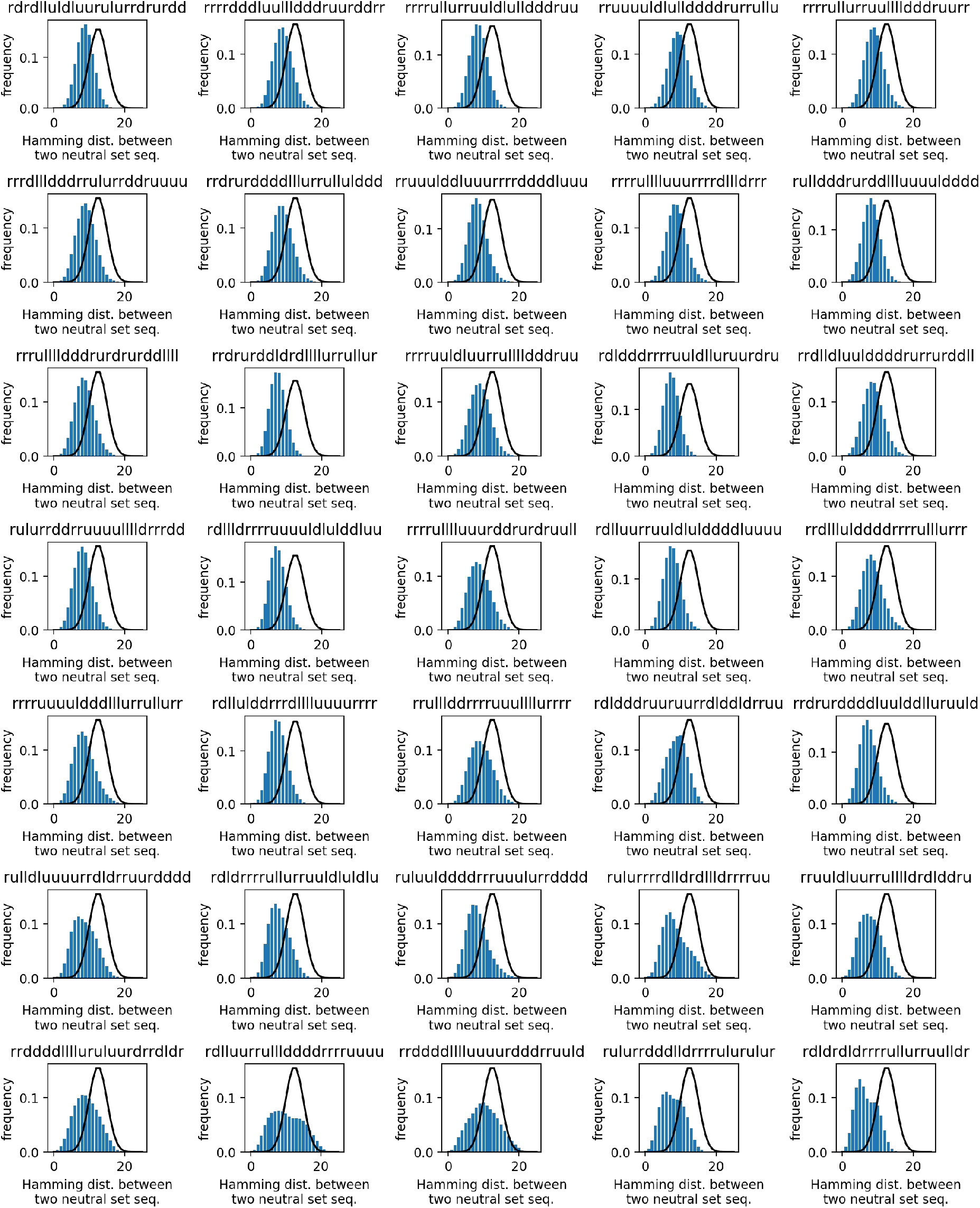
Diversity of sequences within several HP neutral sets: each plot focuses on the neutral set a different initial structure p (using the same selection as the figures above). For each neutral set, we plot the Hamming distance distribution between different sequences in our neutral set (blue histogram), together with the Hamming distance distribution of random sequences (black line), as in our previous work on RNAshapes [2]. This data is based on 10^3^ sequences chosen randomly with replacement from each neutral set.

### S6 Different reduced temperature parameters in the HP model

**Figure S19:**
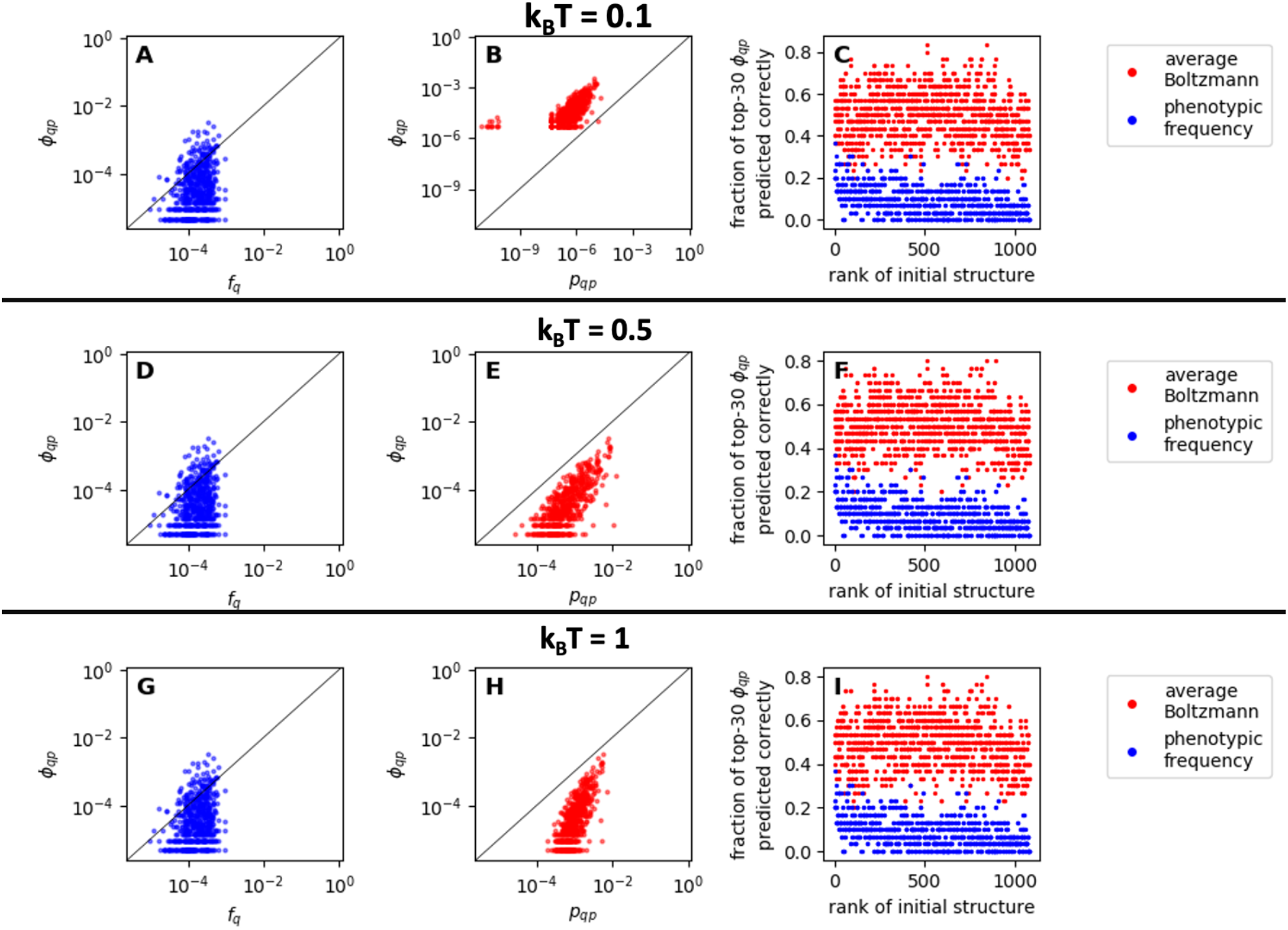
Comparison of the existing and new approach for HP protein structures at different reduced temperatures: this plot is a repetition of the analysis in the main text (Fig. 2D-F), but for three different reduced temperatures, one in each row - k_B_T = 0.1, k_B_T = 0.5 and k_B_T = 1. The example used as an initial structure p in the first two columns is the same as in the main text.

For RNA, the energy model is based on measured parameters that apply at a default temperature of *T* = 37° [21], and thus this is the default temperature for ViennaRNA structure predictions. This means that the temperature we use to calculate Boltzmann weights *w*_*s*_ in the expression *w*_*s*_ */* exp(−*G*_*s*_*/*(*k*_*B*_*T*)) should be set to *T* = 37° for consistency.

The HP protein model is more abstract, the free energy values are dimensionless (for example ref [22]) and thus such a direct link to physical temperatures does not exist. Thus, we need to set a reduced temperature relative to the dimensionless contact energy parameters between *H* and *P* residues (*E*_*HH*_ = *-*1 and *E*_*HP*_ = *E*_*PH*_ = *E*_*PP*_ = 0). In ref [23], a reduced temperature of *k*_*B*_*T* = 1 was used. We investigate three reduced temperatures: *k*_*B*_*T* = 0.1, *k*_*B*_*T* = 0.5 and *k*_*B*_*T* = 1. We find that likely phenotypic transitions are well-captured by our new Boltzmann-frequency-based approach for all three reduced temperatures (Fig. S19). In addition, we find that the correlation between the mean Boltzmann frequency *p*_*q*_ of a structure *q* and its phenotypic frequency *f*_*q*_ holds in all cases, whether minimum-free-energy (mfe) structures are included or not (left and centre columns of Fig. S20). Thus, our qualitative results are robust to changes in the reduced temperature.

**Figure S20:**
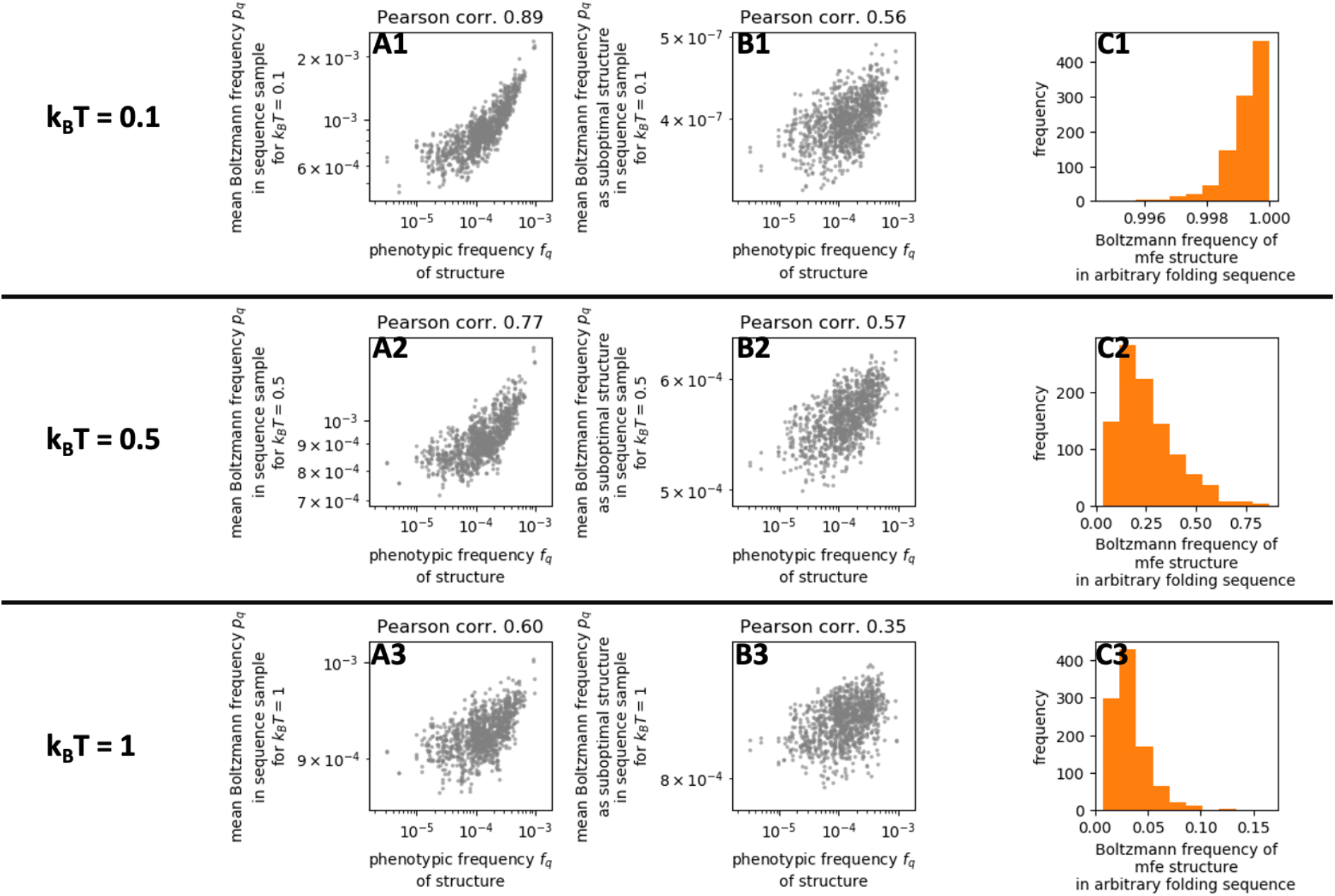
Mean Boltzmann frequencies p_q_ for structure q and typical Boltzmann frequencies at different reduced temperatures: each row uses a different reduced temperature: k_B_T = 0.1, k_B_T = 0.5 and k_B_T = 1 In each row, column A shows the average Boltzmann frequency p_q_ of structure q for random sequences against the phenotypic frequency f_q_ of that structure. We find a positive correlation in all three cases, but this becomes weaker as high-energy states are populated more evenly at higher temperature. Column B shows the same data, except that sequences where the given structure is the mfe structure are excluded from the p_q_ calculations, as in Fig S1. Column C shows the distribution of the Boltzmann frequencies of the mfe structure for 10^3^ random sequences with a unique mfe structure.

While our results hold for all three reduced temperatures, the data for different reduced temperatures shows two qualitative differences: First, the strength of the correlations differs for different temperatures. This is what we might expect since all structures will have equal Boltzmann frequencies as *T* → ∞, whereas the ground states and thus the phenotypic frequencies are unaffected by temperature changes in this model (since entropic contributions are not part of the model). Secondly (right column of Fig. S20), we note that for *k*_*B*_*T* = 0.1, a typical folding sequence folds into its mfe structure with a Boltzmann probability of ⪆ 0.998, whereas for *k*_*B*_*T* = 1, this decreases to ⪅ 0.005. Therefore, we use a value of *k*_*B*_*T* = 0.5 throughout the main text, so that a typical sequence folds primarily into its mfe structure but has some plasticity that corresponds to protein dynamics or even fold-switching as found in ref [24].

## Footnotes

1 Using the expression in ref [17], we estimate there are *≈* 10^6^ structures of length *L* = 30 with no isolated base pairs.

## References

[1] S. F. Greenbury, S. Schaper, S. E. Ahnert, and A. A. Louis, Genetic correlations greatly increase mutational robustness and can both reduce and enhance evolvability, PLOS Computational Biology 12, e1004773 (2016).

[2] A. Wagner, The origins of evolutionary innovations: a theory of transformative change in living systems (OUP Oxford, 2011) Chap. Phenotypic plasticity and innovation.

[3] S. Schaper and A. A. Louis, The arrival of the frequent: how bias in genotype-phenotype maps can steer populations to local optima, PLOS ONE 9, e86635 (2014).

[4] M. C. Cowperthwaite, E. P. Economo, W. R. Har-combe, E. L. Miller, and L. A. Meyers, The ascent of the abundant: how mutational networks constrain evolution, PLOS Computational Biology 4, e1000110 (2008).

[5] M. Stich and S. C. Manrubia, Motif frequency and evolutionary search times in RNA populations, Journal of Theoretical Biology 280, 117 (2011).

[6] K. Dingle, F. Ghaddar, P. Šulc, and A. A. Louis, Phenotype bias determines how natural RNA structures occupy the morphospace of all possible shapes, Molecular biology and evolution 39, msab280 (2022).

[7] K. Dingle, S. Schaper, and A. A. Louis, The structure of the genotype–phenotype map strongly constrains the evolution of non-coding RNA, Interface focus 5, 20150053

[8] S. Manrubia, J. A. Cuesta, J. Aguirre, S. E. Ahnert, L. Altenberg, A. V. Cano, P. Catalán, R. Diaz-Uriarte, S. F. Elena, J. A. García-Martín, P. Hogeweg, B. S. Kha-tri, J. Krug, A. A. Louis, N. S. Martin, J. L. Payne, M. J. Tarnowski, and M. Weiß, From genotypes to organisms: State-of-the-art and perspectives of a cornerstone in evolutionary dynamics, Physics of Life Reviews 38, 55 (2021).

[9] R. Lorenz, S. H. Bernhart, C. H. Zu Siederdissen, H. Tafer, C. Flamm, P. F. Stadler, and I. L. Hofacker, Vi-ennaRNA Package 2.0, Algorithms for molecular biology 6, 26 (2011).

[10] H. Li, C. Tang, and N. S. Wingreen, Emergence of preferred structures in a simple model of protein folding, Science 273, 666 (1996).

[11] B. M. Stadler, P. F. Stadler, G. P. Wagner, and W. Fontana, The topology of the possible: Formal spaces underlying patterns of evolutionary change, Journal of Theoretical Biology 213, 241 (2001).

[12] W. Fontana and P. Schuster, Continuity in evolution: on the nature of transitions, Science 280, 1451 (1998).

[13] J. Vīksna and D. Gilbert, Assessment of the probabilities for evolutionary structural changes in protein folds, Bioinformatics 23, 832 (2007).

[14] I. Coluzza, J. T. MacDonald, M. I. Sadowski, W. R. Taylor, and R. A. Goldstein, Analytic markovian rates for generalized protein structure evolution, PloS one 7, e34228 (2012).

[15] K. Dingle, J. K. Novev, S. E. Ahnert, and A. A. Louis, Predicting phenotype transition probabilities via conditional algorithmic probability approximations, Journal of the Royal Society Interface 19, 20220694 (2022).

[16] S. Wuchty, W. Fontana, I. L. Hofacker, and P. Schuster, Complete suboptimal folding of RNA and the stability of secondary structures, Biopolymers: Original Research on Biomolecules 49, 145 (1999).

[17] L. W. Ancel and W. Fontana, Plasticity, evolvability, and modularity in RNA, Journal of Experimental Zoology 288, 242 (2000).

[18] I. Derényi and G. J. Szöllősi, Effective Temperature of Mutations, Physical Review Letters 114, 058101 (2015).

[19] Q.-Y. Tang and K. Kaneko, Dynamics-evolution correspondence in protein structures, Physical Review Letters 127, 098103 (2021).

[20] A. Wagner, Mutational robustness accelerates the origin of novel RNA phenotypes through phenotypic plasticity, Biophysical journal 106, 955 (2014).

[21] N. S. Martin and S. E. Ahnert, Insertions and deletions in the RNA sequence–structure map, Journal of the Royal Society Interface 18, 20210380 (2021).

[22] N. S. Martin and S. E. Ahnert, Fast free-energy-based neutral set size estimates for the RNA genotype– phenotype map, Journal of the Royal Society Interface 19, 20220072 (2022).

[23] A. V. Finkelstein, A. Y. Badretdinov, and A. M. Gutin, Why do protein architectures have Boltzmann-like statistics?, Proteins: Structure, Function, and Bioinformatics 23, 142 (1995).

[24] H. Li, C. Tang, and N. S. Wingreen, Are protein folds atypical?, Proceedings of the National Academy of Sciences 95, 4987 (1998).

[25] E. I. Shakhnovich and A. M. Gutin, Engineering of stable and fast-folding sequences of model proteins., Proceedings of the National Academy of Sciences 90, 7195 (1993).

[26] A. Wagner, Robustness and evolvability: a paradox resolved, Proceedings of the Royal Society B: Biological Sciences 275, 91 (2008).

[27] K. Sato, Y. Ito, T. Yomo, and K. Kaneko, On the relation between fluctuation and response in biological systems, Proceedings of the National Academy of Sciences 100, 14086 (2003).

[28] C. Furusawa and K. Kaneko, Global relationships in fluctuation and response in adaptive evolution, Journal of The Royal Society Interface 12, 20150482 (2015).

[29] C. Espinosa-Soto, O. C. Martin, and A. Wagner, Phenotypic plasticity can facilitate adaptive evolution in gene regulatory circuits, BMC evolutionary biology 11, 1 (2011).

[30] D. S. Tawfik, Messy biology and the origins of evolutionary innovations, Nature chemical biology 6, 692 (2010).

[31] A. Leo-Macias, P. Lopez-Romero, D. Lupyan, D. Zerbino, and A. R. Ortiz, An analysis of core deformations in protein superfamilies, Biophysical journal 88, 1291 (2005).

[32] C. Li, W. Qian, C. J. Maclean, and J. Zhang, The fitness landscape of a tRNA gene, Science 352, 837 (2016).

[33] R. Wroe, H. S. Chan, and E. Bornberg-Bauer, A structural model of latent evolutionary potentials underlying neutral networks in proteins, HFSP journal 1, 79 (2007).

[34] T. Jörg, O. C. Martin, and A. Wagner, Neutral network sizes of biological RNA molecules can be computed and are not atypically small, BMC Bioinformatics 9, 464 (2008).

[35] M. Weiß and S. E. Ahnert, Using small samples to estimate neutral component size and robustness in the genotype–phenotype map of RNA secondary structure, Journal of the Royal Society Interface 17, 20190784 (2020).

[36] K. Dingle, C. Q. Camargo, and A. A. Louis, Input– output maps are strongly biased towards simple outputs, Nature communications 9, 761 (2018).

[37] S. F. Greenbury, General properties of genotype-phenotype maps for biological self-assembly, Ph.D. thesis, University of Cambridge (2014).

[38] D. H. Turner and D. H. Mathews, NNDB: the nearest neighbor parameter database for predicting stability of nucleic acid secondary structure, Nucleic Acids Research 38, D280 (2009).

[39] R. Rezazadegan, C. Barrett, and C. Reidys, Multiplicity of phenotypes and RNA evolution, Journal of theoretical biology 447, 139 (2018).

[40] N. Martin and S. Ahnert, Thermodynamics and neutral sets in the RNA sequence-structure map, Europhysics Letters 139, 37001 (2022).

[41] A. Irbäck and C. Troein, Enumerating designing sequences in the hp model, Journal of Biological Physics 28, 1 (2002).

[42] S. F. Greenbury, A. A. Louis, and S. E. Ahnert, The structure of genotype-phenotype maps makes fitness landscapes navigable, Nature Ecology & Evolution 6, 1742 (2022).

[43] E. Bornberg-Bauer, How are model protein structures distributed in sequence space?, Biophysical journal 73, 2393 (1997).

## References

1 M. Weiß and S. E. Ahnert, “Using small samples to estimate neutral component size and robustness in the genotype–phenotype map of RNA secondary structure”, Journal of the Royal Society Interface 17, 20190784 (2020).

2 N. S. Martin and S. E. Ahnert, “Insertions and deletions in the RNA sequence–structure map”, Journal of the Royal Society Interface 18, 20210380 (2021).

3 S. Schaper and A. A. Louis, “The arrival of the frequent: how bias in genotype-phenotype maps can steer populations to local optima”, PLOS ONE 9, e86635 (2014).

4 S. F. Greenbury, S. Schaper, S. E. Ahnert, and A. A. Louis, “Genetic correlations greatly increase mutational robustness and can both reduce and enhance evolvability”, PLOS Computational Biology 12, e1004773 (2016).

5 R. Lorenz, S. H. Bernhart, C. H. Zu Siederdissen, H. Tafer, C. Flamm, P. F. Stadler, and I. L. Hofacker, “ViennaRNA Package 2.0”, Algorithms for molecular biology 6, 26 (2011).

6 R. Wroe, H. S. Chan, and E. Bornberg-Bauer, “A structural model of latent evolutionary potentials underlying neutral networks in proteins”, HFSP journal 1, 79 (2007).

7 I. L. Hofacker, “RNA secondary structure analysis using the Vienna RNA Package”, Current Protocols in Bioinformatics 4, 12.2.1–12.2.12 (2003).

8 A. Godzik, J. Skolnick, and A. Kolinski, “Regularities in interaction patterns of globular proteins”, Protein Engineering, Design and Selection 6, 801–810 (1993).

9 H. Li, C. Tang, and N. S. Wingreen, “Are protein folds atypical?”, Proceedings of the National Academy of Sciences 95, 4987–4990 (1998).

10 B. M. Stadler, P. F. Stadler, G. P. Wagner, and W. Fontana, “The topology of the possible: formal spaces underlying patterns of evolutionary change”, Journal of Theoretical Biology 213, 241–274 (2001).

11 W. Fontana and P. Schuster, “Continuity in evolution: on the nature of transitions”, Science 280, 1451–1455 (1998).

12 G. Steger and R. Giegerich, “14. RNA structure prediction”, in RNA Structure and Folding, edited by D. Klostermeier and C. Hammann (De Gruyter, 2013), pp. 335–362.

13 N. S. Martin and S. E. Ahnert, “Fast free-energy-based neutral set size estimates for the RNA genotype–phenotype map”, Journal of the Royal Society Interface 19, 20220072 (2022).

14 K. Dingle, J. K. Novev, S. E. Ahnert, and A. A. Louis, “Predicting phenotype transition probabilities via conditional algorithmic probability approximations”, Journal of the Royal Society Interface 19, 20220694 (2022).

15 K. Dingle, C. Q. Camargo, and A. A. Louis, “Input–output maps are strongly biased towards simple outputs”, Nature communications 9, 761 (2018).

16 I. G. Johnston, K. Dingle, S. F. Greenbury, C. Q. Camargo, J. P. Doye, S. E. Ahnert, and A. A. Louis, “Symmetry and simplicity spontaneously emerge from the algorithmic nature of evolution”, Proceedings of the National Academy of Sciences 119, e2113883119 (2022).

17 M. E. Nebel and A. Scheid, “On quantitative effects of RNA shape abstraction”, Theory in Biosciences 128, 211 (2009).

18 E. Ferrada and A. Wagner, “A comparison of genotype-phenotype maps for RNA and proteins”, Biophysical journal 102, 1916–1925 (2012).

19 U. Bastolla, H. E. Roman, and M. Vendruscolo, “Neutral evolution of model proteins: diffusion in sequence space and overdispersion”, Journal of theoretical biology 200, 49–64 (1999).

20 P. Schuster, W. Fontana, P. F. Stadler, and I. L. Hofacker, “From sequences to shapes and back: a case study in RNA secondary structures”, Proceedings of the Royal Society of London. Series B: Biological Sciences 255, 279–284 (1994).

21 D. H. Mathews and D. H. Turner, “Prediction of rna secondary structure by free energy minimization”, Current opinion in structural biology 16, 270–278 (2006).

22 H. Li, C. Tang, and N. S. Wingreen, “Emergence of preferred structures in a simple model of protein folding”, Science 273, 666–669 (1996).

23 J. D. Bloom, J. J. Silberg, C. O. Wilke, D. A. Drummond, C. Adami, and F. H. Arnold, “Thermodynamic prediction of protein neutrality”, Proceedings of the National Academy of Sciences 102, 606–611 (2005).

24 L. L. Porter and L. L. Looger, “Extant fold-switching proteins are widespread”, Proceedings of the National Academy of Sciences 115, 5968–5973 (2018).

